# Representation of Borders and Swimming Kinematics in the Brain of Freely-Navigating Fish

**DOI:** 10.1101/291013

**Authors:** Ehud Vinepinsky, Lear Cohen, Shay Perchik, Ohad Ben-Shahar, Opher Donchin, Ronen Segev

## Abstract

Like most animals, the survival of fish depends crucially on navigation in space. This capacity has been documented in numerous behavioral studies that have revealed navigation strategies and the sensory modalities used for navigation. However, virtually nothing is known about how freely swimming fish represent space and locomotion in the brain to enable successful navigation. Using a novel wireless neural recording system, we measured the activity of single neurons in the goldfish lateral pallium, a brain region known to be involved in spatial memory and navigation, while the fish swam freely in a two-dimensional water tank. Four cell types were identified: border cells, head direction cells, speed cells and conjunction head direction with speed. Border cells were active when the fish was near the boundary of the environment. Head direction cells were shown to encode head direction. Speed cells only encoded the absolute speed independent of direction suggestive of an odometry signal. Finally, the conjunction of head direction with speed cells represented the velocity of the fish. This study thus sheds light on how information related to navigation is represented in the brain of swimming fish, and addresses the fundamental question of the neural basis of navigation in this diverse group of vertebrates. The similarities between our observations in fish and earlier findings in mammals may indicate that the networks controlling navigation in vertebrate originate from an ancient circuit common across vertebrates.

**Summary:** Navigation is a fundamental behavioral capacity facilitating survival in many animal species. Fish is one lineage where navigation has been explored behaviorally, but it remains unclear how freely swimming fish represent space and locomotion in the brain. This is a key open question in our understanding of navigation in fish and more generally in understanding the evolutionary origin of the brain’s navigation system. To address this issue, we recorded neuronal signals from the brain of freely swimming goldfish and successfully identified representations of border and swimming kinematics in a brain region known to be associated with navigation. Our findings thus provide a glimpse into the building blocks of the neural representation underlying fish navigation. The similarity of the representation in fish with that of mammals may be key evidence supporting a preserved ancient mechanism across brain evolution.

## Introduction

Navigation is a fundamental behavioral capacity facilitating survival in many animal species [1–4]. It involves the continuous estimation and representation of the agent’s position and direction in the environment, which are implemented in the planning and execution of movements and trajectories towards target locations [5,6]. Navigation has been investigated extensively on numerous taxa across the animal kingdom but attempts to probe its neural substrate have mainly been focused on mammals [7] and insects [8]. In mammals, neurons in the hippocampal formation encode information about the position and orientation of the animal in space [5–7,9,10]. These cells include place cells [11], grid cells [12], head-direction cells [13,14] and other cell types [15,16]. In insects, a ring-shaped neural network in the central complex of the fruit fly was shown to represent its heading direction [8].

To better understand space representation in other taxa, we explored the neural substrate of navigation in the goldfish (*Carassius auratus*). These fish are known to be able to navigate by exploiting either an allocentric or an egocentric frame of reference. This may imply that the goldfish has the ability to build an internal representation of space in the form of a cognitive map [17]. This would include cognitive map-like navigation strategies to find a goal when starting from an unfamiliar initial position, or taking shorter alternative routes (shortcuts) when possible [17–20]. Furthermore, goldfish are known to use many environmental cues, such that any single cue would not be crucial in itself to navigating in the environment [19].

In addition to these behavioral studies, lesion studies in goldfish have shown that the telencephalon is crucial for spatial navigation. A lesion in the lateral pallium in the telencephalon leads to dramatic impairment in allocentric spatial memory and learning, but not when the lesion affects other parts of the telencephalon [19]. These findings are similar to results from lesions studies of the hippocampus in mammals and further strengthen the notion that the lateral pallium in goldfish is a possible homologue of the mammalian hippocampus [17,21] (but see also [22]). While these works have contributed to suggesting where the possible navigation mechanism is located in the brain, they have not addressed the nature of the representation of this information, which is the goal of this study.

## Results

To better understand how facets of space and locomotion are represented in the teleost brain, we measured single cell activity in the lateral pallium of freely behaving goldfish while they explored a water tank. We first trained the fish (13-15 cm body length) to swim in a tank measuring 0.6 m X 0.6 m X 0.2 m (Figure 1A). After the fish became familiar with the water tank and learned to explore its entire environment naturally, we installed a recording system on their heads [23]. We then let a goldfish swim freely, while a camera positioned over the water tank recorded the fish’s location and head orientations. A data logger placed in a waterproof case mounted on the fish’s skull recorded the activity of single cells using tetrodes (Figure 1, see Methods).

**Figure 1.**
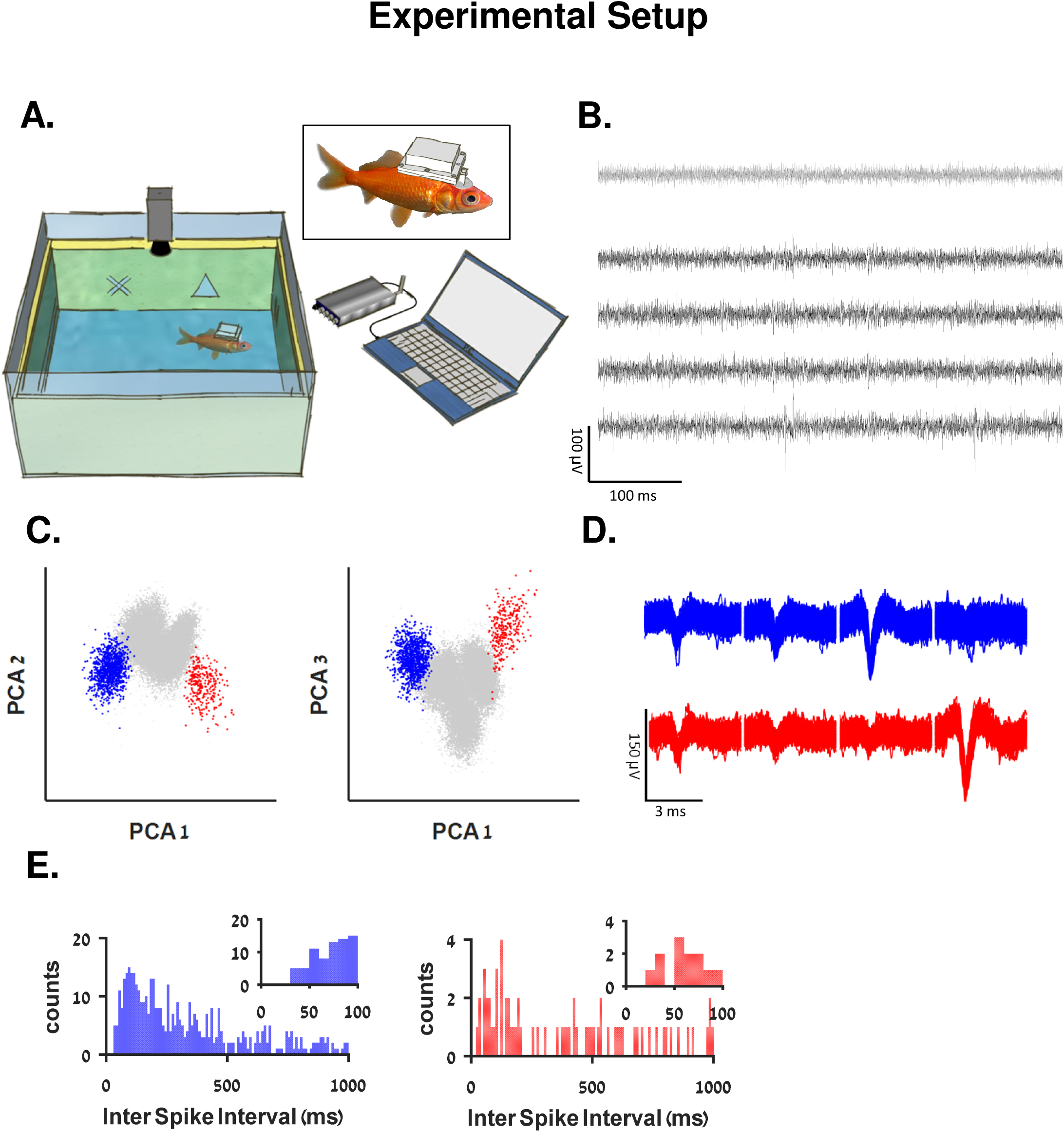
The experimental system. **A.** Schematic overview of the experimental setup: a fish swims freely in a water tank with electrodes and a wireless data logger mounted on its head. The fish’s movements are recorded by a video camera positioned above the tank. The tank’s walls were coated with yellow foam sheets, marked with different visual landmarks. **B.** Example of a raw recording from a tetrode (black traces) and a reference electrode (gray) in the fish’s lateral pallium. Neural activity can be seen in the tetrode alone. **C**. Projection on the first three principal components of the data from the tetrode of all spike candidates that crossed the threshold. **D.** Waveforms of two neurons after spike sorting. Other clusters were not distinguishable from other multiunit activity and neural noise. The blue cluster forms a velocity cell (supplementary Figure 13 A-F) and the red cluster forms a head direction cell (additional examples can be found in supplementary Figure 1). **E.** Inter spike interval histogram of the detected clusters. Insets show there were no violations of the refractory period.

Using this methodology, we identified four types of cells that may constitute the primitives of the goldfish navigation system: border cells, head direction cells, speed cells, and conjunction head direction and speed cells. Border cells were active when the fish was in close proximity to the borders of its environment (i.e., in our case, near the walls of the water tank). Head direction cells were active when the fish’s head was in a specific orientation. Speed cells activity was correlated with the fish’s swimming speed, regardless of the swimming direction. Velocity cells were more active when the fish swam in a particular direction and speed, thus conjugating head direction and speed characteristics.

Examination of the neural activity (red dots, Figure 2A) of a border cell over the fish’s trajectory (black curve) reveals a clear pattern suggesting that this neuron was mainly active when the fish swam near the boundaries of the water tank. This was also documented by the heat map of that neuron (Figure 2B), color-coded from dark blue (zero firing rate) to dark red (maximal firing rate, indicated at the top right side of the panel). To test statistically whether a neuron was a border cell, we first found the minimal distance from the wall in which 75% of the spikes occurred. Then, we defined this distance as the border activity layer (red arrow, Figure 2D), and after implementing a standard shuffling procedure we obtained 5000 shuffled spike trains and measured their border activity layer (Figure 2D). When activity of the cell was significantly closer to the environmental borders (p<0.05, see Methods), the cell was classified as a border cell. As can be seen in the graph of spike distribution vs. distance from the borders (Figure 2C), this result did not depend exclusively on the selection of the distance from the border. Additional examples of border cells are depicted in Figure 2 E-L and in supplementary Figure 3.

**Figure 2.**
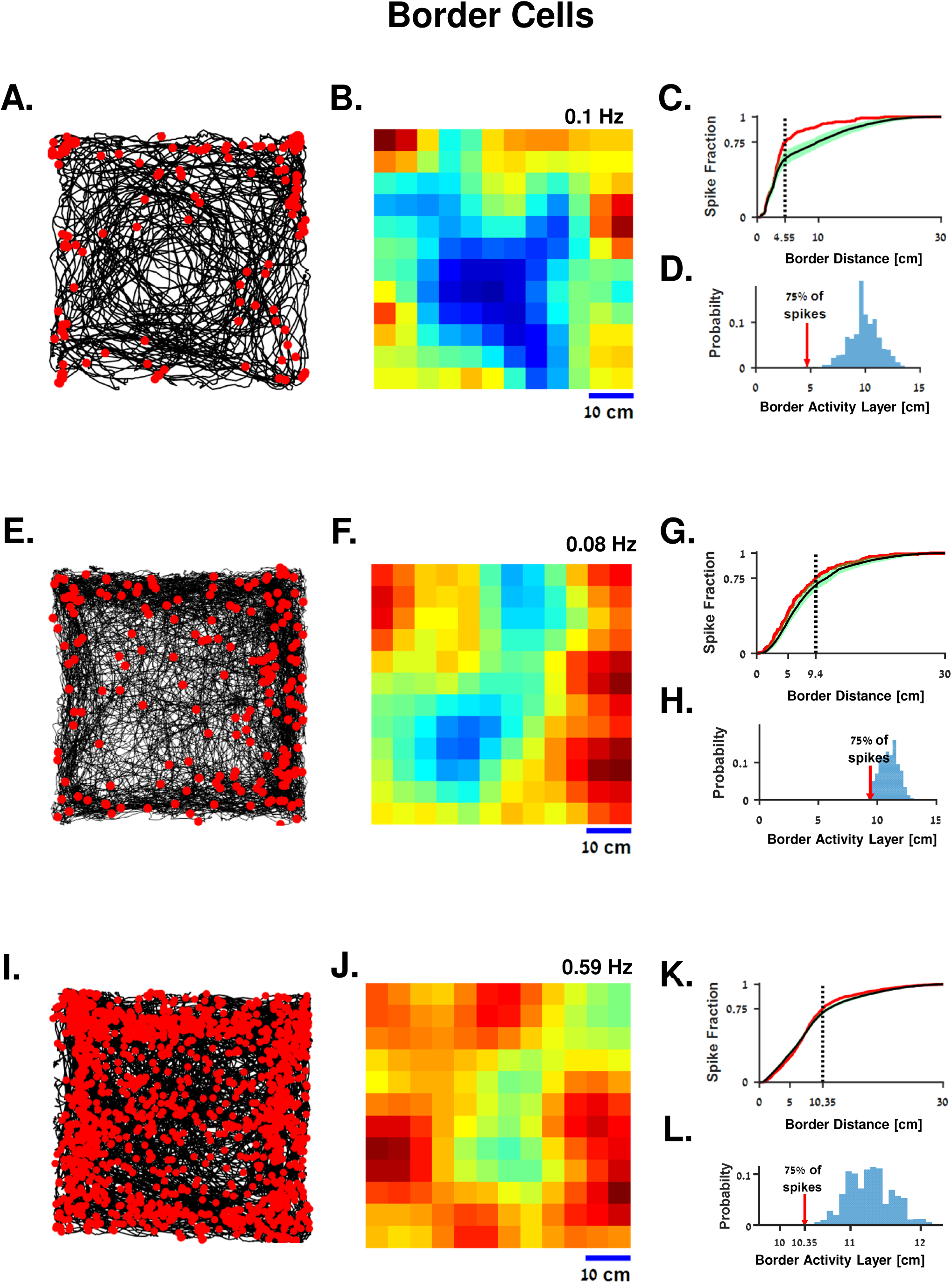
Border encoding cells in the goldfish lateral pallium. **A.** An example of a border cell. The fish trajectory (black curve) is presented together with the location of the fish when each spike of a single cell occurred (red dots). The apparent pattern shows that this neuron was mainly active when the fish was at the boundaries of the tank. Waveform of the cell spiking activity is presented in supplementary Figure 1C. **B.** Firing rate map of the cell in A. The highest spiking probabilities are concentrated at the boundaries. **C and D.** Statistical analysis of firing near the border of the water tank. Border activity layer of shuffled spike trains (black and green curves in panel C represent the mean and 95% confidence interval, respectively) are contrasted with the border activity layer of the cell (red curve in panel C). Panel D indicates the distance from the border in which 75% of spikes fired by the recorded neuron (red arrow) compared to the shuffled data (blue histogram). **E-L.** Another border cell example (additional examples are presented in supplementary Figure 3).

In order to test the stability of the border tuning across trials, and to verify that the border cells indeed encoded the presence of the fish near the arena walls rather than some specific feature of the visual scene, we performed two transfer experiments. In the first experiment, we rotated the environment by 180 degrees. After this manipulation, cell activity was still tuned to the borders of the arena, but did not completely rotate (supplementary Figure 4 A-H). In the second control experiment, we transferred the fish to a circular arena. In this case, the cell still encoded the presence of the fish near the border (supplementary Figure 4 I-P). In a stability over time test, in which the analysis of a one-hour session was split into two halves (supplementary Figure 5), the activity heat map and the border tuning were found to be similar in both parts of the experiment. Finally, population analysis showed that 14 out of 142; i.e., about 10 % of all the well-isolated units recorded in 18 fish, could be classified as border cells.

Head direction cells are neurons which encode the head direction of the fish by generating a higher firing rate in a preferred orientation. Several examples of head direction and non-head direction cells are depicted in Figure 3. Examination of the firing rate as a function head direction revealed the directional tuning of head direction cells (Figure 3 B, E, H, together with the activity across space in panels A, D, G) and quasi-isotropic directional tuning of other non-head direction cells (Figure 3 K and N, together with the activity across space in panels J and M). To test whether a neuron was directionally tuned, we used the standard length of the Rayleigh vector as the head direction index [24]. Then, we employed a standard shuffling procedure to obtain shuffled spike trains and measured the head direction index (also referred to as the score) of each of the shuffled spike trains to obtain a confidence interval. Cells whose head direction index was significantly larger than the shuffled data (p<0.05) were classified as head direction cells (see Figure 3 C, F and I for head direction cell examples, and L and O for non-direction cells examples, see Methods). Population scores are presented in Figure 3P. Additional examples of head direction cells are presented in supplementary Figure 6.

**Figure 3.**
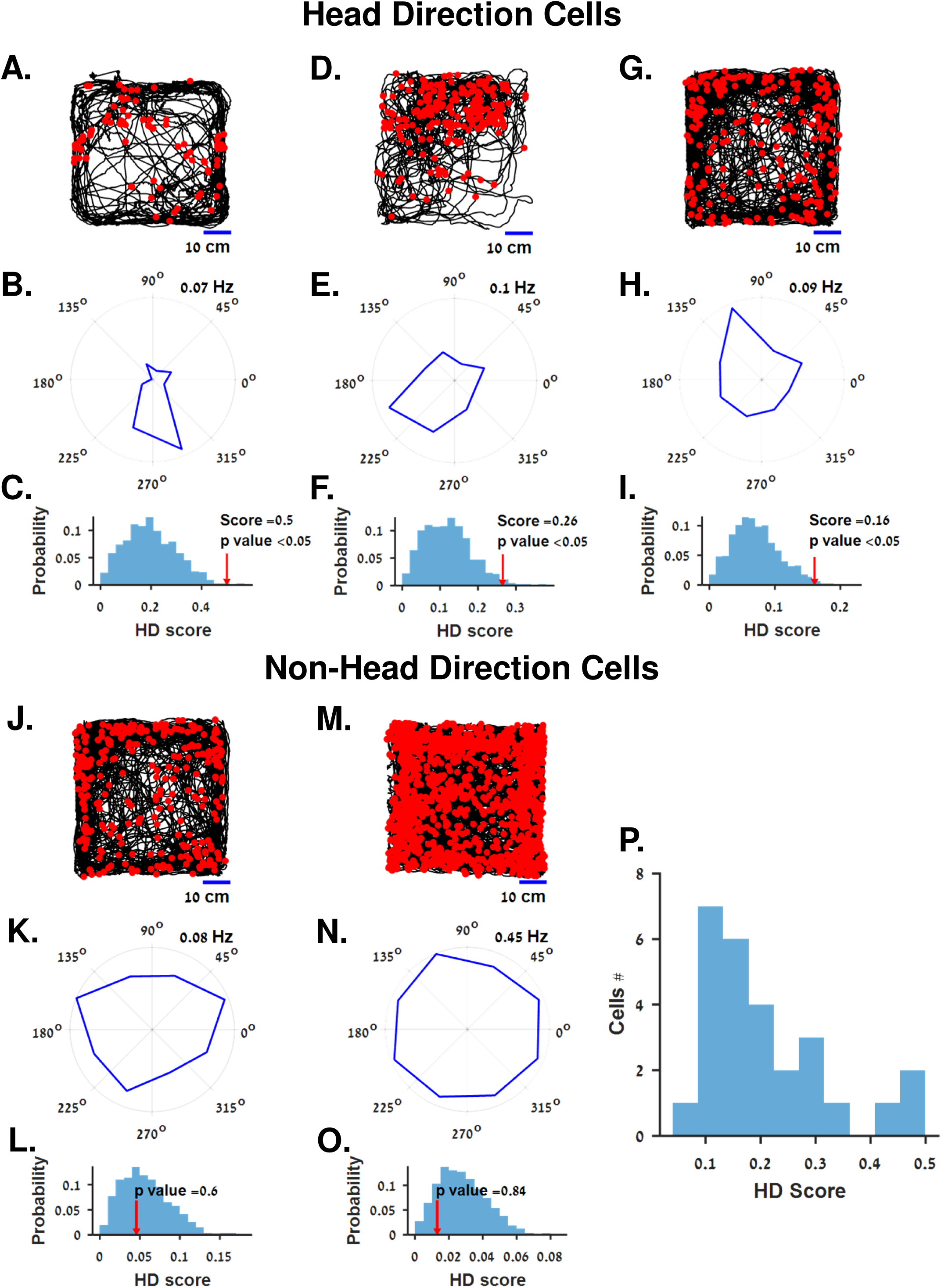
Head direction cells in the goldfish lateral pallium. **A.** The fish’s trajectory (black curve) is presented together with the location of the fish when each spike of a single cell occurred (red dots). **B.** Rate as function of direction for the cell in A. The cellular response is tuned to the negative y-direction. **C.** Statistical analysis of the cell in A and B. The head direction score of the cell is compared with 5000 shuffled spike trains. **D-I.** Two additional examples of head direction cells. **J-O.** Two example of non-head direction cells. **P.** Head direction scores for all head direction cells (N=27).

To test the stability of directional tuning across trials and to determine whether the head direction cells were influenced by internal cues alone, or also by the visual scene, we rotated the water tank with the visual cues by 180 degrees between two recording sessions of head direction cells. After the rotation, the preferred direction of the cell rotated almost 180 degrees (supplementary Figure 7). This may be an indication that the cell was directionally tuned across different environments. In addition, since the rotation was not equal to 180 degrees, it may be the case that the head direction signal could be generated by integrating both external (e.g. visual or olfactory from the body of water) and internal (e.g. vestibular) cues. In total, population analysis identified 27 head direction cells out of 142 well-isolated cells; i.e., 19% of all cells recorded in 18 fish.

The speed cells were tuned to the absolute value of the fish’s swimming velocity and thus represent locomotion in an egocentric coordinate system making them a potential odometry signal. Two examples of speed cells are presented in Figure 4 A-F. The firing rate map in the velocity space (Figure 4 A and D) revealed a pattern showing a tuning to speed, where the firing rate increased with speed (Figure 4 B and E) regardless of the direction of motion (Figure 4 C and F). The correlation coefficient between the neuronal firing rate and fish’s speed was calculated, and a standard shuffling procedure was used to test for speed selectivity of the neurons (see Methods). The correlation coefficient between firing rate and speed showed a statistically significant pattern (insets, Figure 4 B and E). Additional examples are presented in supplementary Figure 8. Fish trajectories and spiking activity associated with all speed cells examples are presented in supplementary Figure 9. Detailed analyses of the distribution of speed across different areas in the water tank; i.e., center vs. border vicinity, showed no apparent differences (supplementary Figure 10). The distribution of maximal speed and speed tuning curve slopes are present in supplementary Figure 10 (I-J). Finally, population analysis indicated that 81 out of 142; i.e., 57% of all well-isolated units recorded in 18 fish, could be classified as speed cells.

**Figure 4.**
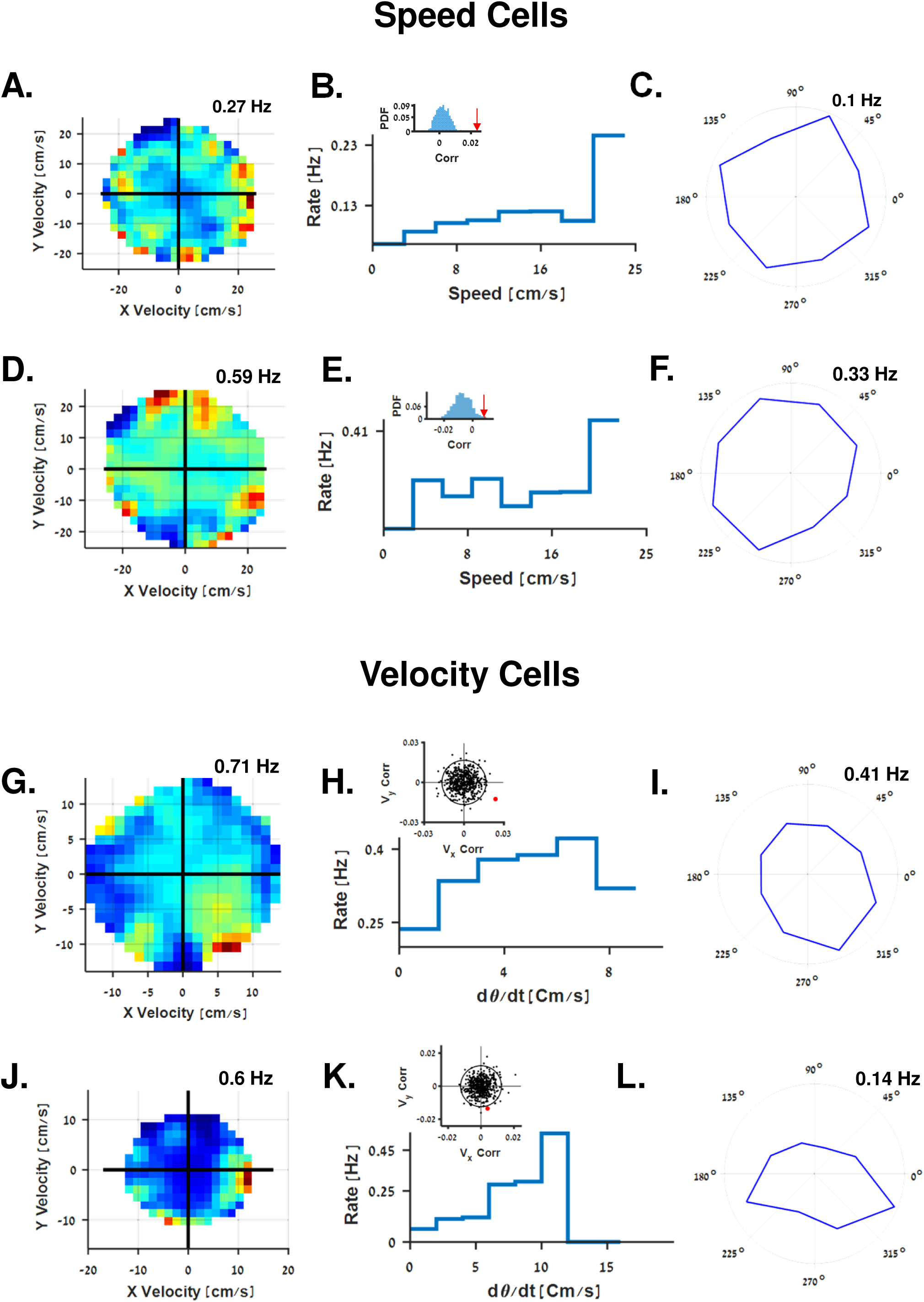
Velocity and speed cells in the goldfish lateral pallium. **A. Example of a speed cell.** Firing rate of the cell as a function of the velocity in the plane. The cell’s tuning to speed is manifested by the fact that the firing rate at the axis origin is low compared to the boundaries of the circle. **B.** Firing rate of the cell in A as a function of the fish’s speed. The firing rate increases with the speed. Inset shows the correlation coefficient between firing rate and speed for the cell in A (red arrow) vs. the correlation coefficient of the firing rate and the speed of 5000 shuffled spike trains. **C.** Rate as a function of head direction for the cell in A. Low score and quasi isotropic shape suggests speed cells are independent of direction preferences. **D-F.** Another example of a speed cell. **G.** An example of a velocity cell. Firing rate of the cell as a function of velocity in the plane. The cell is tuned to the velocity in the positive y-direction. **H.** Rate as a function of velocity in the preferred direction (indicated as 8) of the cell in G. Firing rate increases with velocity. Insets are two-dimensional correlations between firing rate and fish velocity of the cell in G (red dot), compared to those of 5000 shuffled spike trains (black dots. Black circle marks the 95^th^ percentile of the shuffled correlations). **I.** Rate as a function of head direction for the cell in G. Preferred direction corresponds to the preferred swimming direction in G. **J-L.** Another example of a velocity cell.

Conjunction head direction and speed cells are active when the fish swims in a specific direction and speed, regardless of position; i.e., these cells encode velocity. Velocity cells fire is correlated with a preferred direction and swimming speed of the fish. Hence, this cell type represents locomotion in space in an allocentric coordinate; i.e., world centered reference frame. Examples of velocity cells are presented in Figure 4 G-L. The color-coded firing rate map in the velocity space (Figure 4 G and J) showed that cellular activity was correlated with both direction and speed. Similar patterns also emerged from the head direction preference of these cells (Figure 4 I and L), and corresponded to the firing rate as a function of speed in the preferred direction (Figure 4 H and K). To test statistically whether a neuron was a velocity cell, we employed a standard shuffling procedure, and compared the two-dimensional correlation between firing rate and velocity of the recorded cell with the corresponding correlations of the shuffled neurons (Insets, Figure 4 H and K, p<0.05, see Methods). Additional examples are presented in supplementary Figure 11. The fish trajectories and spiking activity associated with all velocity cell examples are presented in supplementary Figure 12.

To test the velocity tuning across recording sessions, we transferred the fish from a square water tank into a circular one, while recording the same velocity cell. In both sessions, the cell was classified as a velocity encoding unit, but the preferred direction changed (supplementary Figure 13 A-F). A test of the stability of the velocity cell indicated no difference between the first and second half of the same recording session (supplementary Figure 13 G-L). In total, 25 out of 142; i.e., 17% of all units recorded in 18 fish could be classified as velocity cells.

In addition, we located the recording site using standard post-recording anatomical procedures in ten of the fish (Figure 5, more details in supplementary Figure 14, see Methods). The different cell types were found mainly in the Dlv, Dp and Dd regions, but the different brain structures could not be differentiated in terms of their functionality. Note that the division between the Dlv and Dp is subject to ongoing debate [25–27].

**Figure 5.**
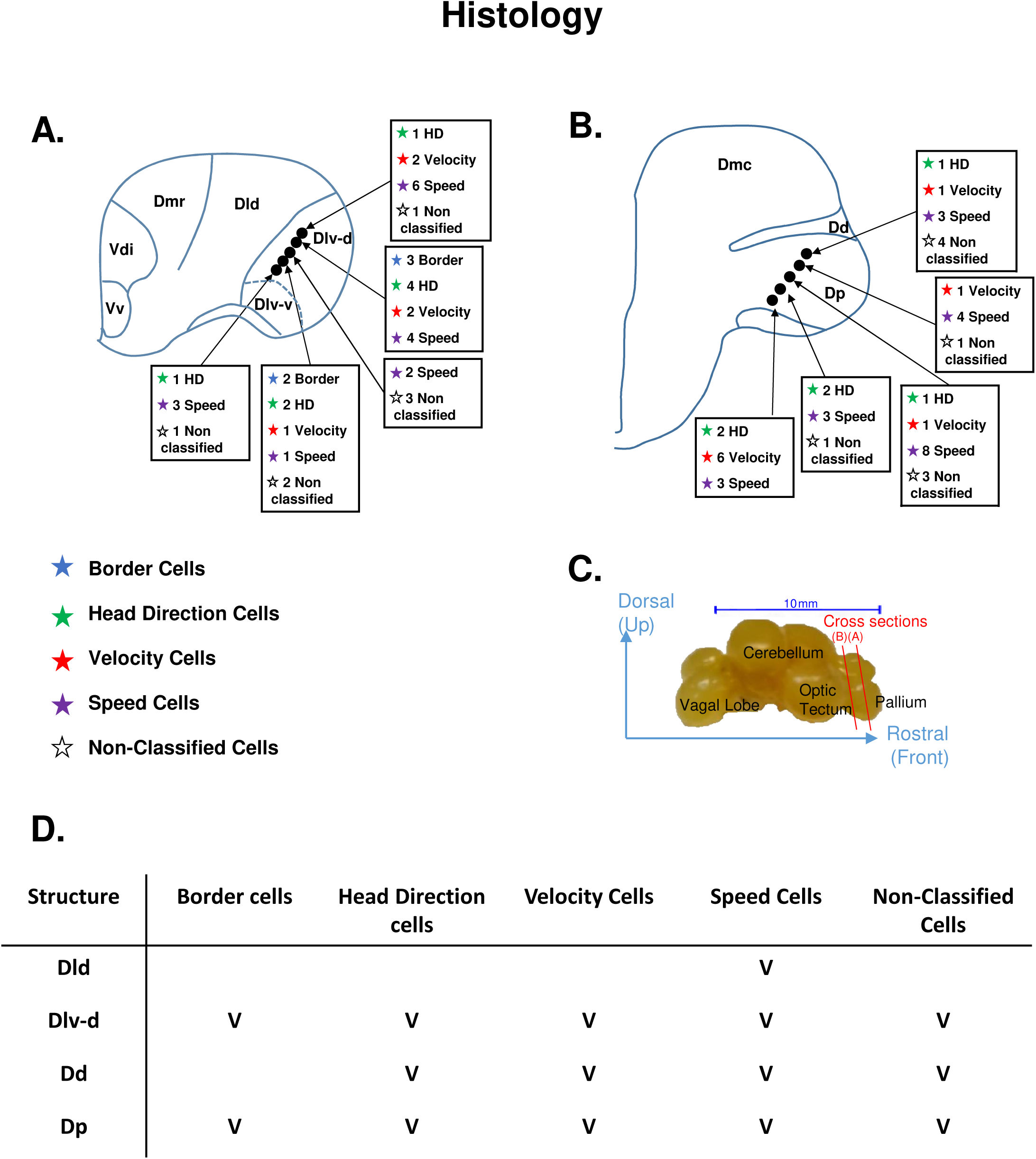
Distribution of cells in the goldfish brain. **A-B**. Two examples of recording locations and cell types found in those locations (anatomical diagram based on Northchutt [25]). **C**. Fish’s brain structures. Red cross-sections A and B correspond to the brain sections presented in panels A and B, respectively. **D.** Summary of the locations in 10 fish and the type of cells found in these locations.

## Discussion

We found border-like, head direction, velocity and speed cells that represent different components of position and locomotion in the goldfish brain. These neurons were found mainly in the Dlv/Dp and Dd structures in the goldfish pallium, which suggests these regions contain a variety of neurons that represent features of space, direction and speed. These cell types represent the minimal requirements of a navigation system; namely, a compass, a reference frame and an odometer. The head direction cells provide a continuous directional signal which the brain can use as a compass [28]. The border cells can provide a reference frame [29] and integration over time of speed cells can provide information about distance travelled. Note that although the goldfish border-like cells differ from the mammalian border cells by their tuning to more than one wall of the arena, they still fulfil the same function of providing a reference frame.

These findings hint that these cell types may constitute the basic building blocks of the goldfish navigation system and can thus shed new light on theories of navigation systems derived from observations of the mammalian system alone. Further investigations are needed to obtain a complete functional map of this region and to understand whether it plays a foundational role in the neural navigation system of the teleost brain.

Finding cell types similar to the ones found in the mammalian hippocampal formation in the fish’s lateral pallium, in the form of border-like cells, head direction cells, speed cells and velocity cells suggests that the representation of borders and self-kinematics may have evolved from an ancient neural circuit common to both teleost and mammals. Therefore, these results have important implications for current theories on the evolution of brain function in the teleost and outside the teleost lineage. They suggest that the actual homologue of the lateral pallium is the hippocampal formation, which is consisted one of several sub-regions in the mammalian brain, including the hippocampus, the entorhinal cortex, the subiculum and the post-subiculum [30]. Currently, however, it is impossible to determine the exact homology between the divisions of the mammalian hippocampal formation and the divisions of the pallium in the teleost brain. In fact, it remains unclear whether there is such a division. Given the disparity between what is known about connectivity maps in the two lineages, fish brain connectivity is poorly understood compared to the relatively well-known connectivity of the mammalian brain [30].

Although there is a robust consensus regarding the inventory of cells that represent space in the mammalian brain [5,7,10], there is no accepted theory based on empirical data as to how these spatial representation components are integrated in the brain into a functioning navigation system [5]. One solution can emerge from studying the neural mechanisms underlying navigation in other vertebrate lineages, since elucidating the mechanisms of space representation in different vertebrate taxa can help decipher how the elementary building blocks of navigation are integrated into a functional navigation system [31–35]. Crucially, comparative studies can help determine whether the mammalian navigation system is unique or an instance of a more general biological design. Thus, a comparative approach may help resolve the key question of the critical components making up a functioning navigation system.

Only small number of studies have attempted to study the neural representation of self-kinematic and space outside of the mammalian linage. Bingman et al. [36] analysis of place cells in the pigeon’s hippocampus homolog only found preliminary evidence of spatial encoding. Canfield et al. published a method paper describing an extracellular recording system in tethered goldfish, and reported preliminary evidence for speed and place encoding, but without supporting statistics. Arhens et al. [37,38] studied the larval zebrafish brain during fictive navigation in virtual environments with inconclusive results. A complementary study in the telencephalon of weakly electric fish observed supporting evidence for cells which are more active near objects and borders [39]. Finally, studies in the fruit fly have suggested it can represent orientation with respect to salient landmarks in the environment [8,40].

Spiking activity in the fish pallium is rather sparse, without the characteristic activity bursts of many spikes found in certain loci in the mammalian hippocampal formation. Thus fish firing rates are much lower. In addition, low firing rates can also be found in spatial encoding cells in the mammalian dentate gyrus [41]. Similar firing rates were recently observed in the electric fish telencephalon [39]. This low firing rate means that the fish modulates location and kinematics at a higher rate than the modulation of spiking activity in our recordings. This may suggest that the goldfish does not represent speed, location or any other spatial information via a single cell. Rather, at each point in time, a population code generates a much more accurate representation of speed modulation at the population level.

Thus overall, this study constitutes one step toward a better understanding of the navigation system in non-mammalian vertebrates. This could shed light on the basic building blocks of this system across all vertebrates.

## Methods

### Animals

All goldfish experiments were approved by the Ben-Gurion University of the Negev Institutional Animal Care and Use Committee and were in accordance with government regulations of the State of Israel. Goldfish (Carassius auratus), 13-15 cm in body length, 80-120 g body weight were used in this study. A total of 18 fish were used for the recordings. The fish were kept in a water tank at room temperature. The room was illuminated with artificial light on a 12/12 h day-night cycle. Fish were kept in the home water tank and were brought to the experimental water tank for recordings.

### Surgery and Stereotaxic

Surgery was done outside of the water while the fish was anesthetized and perfused through its mouth (MS-222 200 mg l−1, NaHCO3 400 mg l−1, Cat A-5040 and Cat S-5761, Sigma-Aldrich, USA) as described by Vinepinsky et al.[23]. The desired brain location was targeted by constituting the anterior midmargin of the posterior commissure as the zero point for the stereotaxic procedure as described by Peter and Gil [42]. From the zero point, using a mechanical manipulator, we moved the tetrodes 1 mm laterally, 1 mm ventrally and 1.5 mm anteriorly.

### Wireless electrophysiology

The behavioral fish electrophysiology is described in detail in Vinepinsky et al.[23]. Briefly, the experimental setup for recording extracellular signals from the brain of freely swimming goldfish is based on a small data logger (Mouselog-16, Deuteron Technologies Ltd., Jerusalem, Israel) connected to an implant on the fish’s skull and receives input from one or two tetrodes that are mounted in the fish brain. In addition, we place a reference electrode for detection of possible motion artifacts. To protect the electronics, the neural logger is placed in a waterproof case (Figure 1A). The data logger is controlled wirelessly by a computer via a transceiver (Deuteron Technologies Ltd., Jerusalem, Israel) and records the neural signals for up to 4 hours per session. The fish’s behavior is monitored continuously with a video camera positioned over the tank. The video is synchronized with the recording by feeding a trigger signal from the logger into the camera. A Styrofoam marker mounted on the box which contains the logger for the entire implant (tetrodes, box, logger and battery) is buoyancy neutral (i.e. total average density of 1 gr/cm^3^). The front and back ends of the Styrofoam marker are painted in different colors to easily determine the swimming direction automatically from the video recordings. A total of 142 neurons in 18 fish were monitored.

To ensure our recordings were free of motion artifacts, at the end of each surgery, while the fish was still under anesthesia, we performed a control experiment where we moved the fish in the water tank while recording electrical activity. This included moving the fish around and bumping its body and electronics box (see example in supplementary Figure 2). Only trials devoid of any motion artifact were used for further analysis. Spike waveforms that were present simultaneously on the two recording tetrodes or in the reference electrode were removed from the analysis.

Spike sorting was done by manual clustering using PCA analysis of the spike amplitudes, widths, and waveforms across all the electrodes in each tetrode. All analyses were conducted using an in-house Matlab program (see examples in Figure 1B-D and in supplementary Figure 1).

### Water tank

The water tank for the experiment was 0.6 m X 0.6 m X 0.2 m in size and coated with a foam sheet (Figure 1A). Visual landmarks were marked on the walls of the foam sheets (supplementary Figure 15). A camera above the water tank was used to localize the fish in the X-Y plane. A circular arena with a radius of 0.34 m and a height of 0.2 m was used for some of the control experiments (supplementary Figure 4 and 13)

### Training

Prior to surgery, the fish were trained to explore the entire water tank since in an unfamiliar environment fish tend to stay near the walls or barely swim. Training involved allowing the fish to swim freely in the tank for 20 minutes a day for several days while an automatic feeder, positioned above the center of the tank, fed the fish as soon as it approached the center of the tank. After about a week of training, most fish were familiar with the water tank and explored it efficiently. After training, the fish underwent surgery to implant the tetrodes and the logger case.

### Camera and synchronization

All sessions were recorded using a “Gopro hero 4 black” camera at 24 FPS, HD resolution and a linear field of view. In order to synchronize the neural activity and the video recordings, we used the synchronization system provided with the Mouselog-16 by Deuteron Technologies. We set the Mouselog-16 transceiver to deliver a 100ms wide pulse at intervals of 10 and 20 seconds. The signal was then sent to the camera’s audio input. The camera’s timing was later adjusted to the Mouselog-16 timing using these pulses.

### Recording sessions

Each recording session included synchronized recordings from the Neurolog-16 and the camera system while the fish navigated freely in the water tank. Recording sessions lasted about 1 hour.

### Analysis

After the experiment, the data from the cameras and the Neurolog-16 were analyzed. This was done by detecting the action potential timings using a standard spike-sorting algorithm [43,44]. Then, the correlation coefficients between the fish’s behavior, location and orientation and the neural activity were calculated to determine how space was represented during the task. Subsequently, the brains of ten of the fish were fixed and sliced to reveal the electrode position in the brain (i.e. the lateral pallium or any other brain area targeted for recording).

### Data analysis: Fish trajectory and firing rate map

The fish’s location and orientation in each video frame was detected using the colored Styrofoam marker that was attached to the case containing the logger. The water tank was tessellated to 5 cm^2^ bins and smoothed using a 2- dimensional Gaussian (sigma = 7.5 cm). This yielded two auxiliary maps which indicated how many spikes occurred in each bin and how much time the fish spent in each bin. The firing rate map for each neuron was obtained by dividing the two maps bin by bin. Bins that were not visited enough by the fish were discarded from the analysis and appear on the map as white bins. Sessions in which the fish only swam around the borders were discarded.

### Data analysis of border cells

To determine which cells could be classified as border cells, we measured the width of the border activity layer near the walls of the water tank which contained 75% of the spikes. We compared this to 5000 shuffled spike trains. Each shuffled spike train was obtained by first calculating the inter-spike intervals, shuffling the inter-spike intervals by random permutation and using the cumulative sum to obtain the shuffled spike train. Then we calculated the probability of observing a width of border activity layer by comparing the value obtained for the recorded cell to the values calculated for the shuffled spike trains. Cells with a p value below 0.05 were considered to be border cells (Figure 2 D, H and L). Sensitivity analysis on the fraction of spikes the activity layer should contain was done by calculating the activity layer which contained 0-100% of the spikes (Figure 2 C, G and K).

### Data analysis of head direction cells

The head direction score was defined as the mean Rayleigh vector length on the tuning curve as is common practice [24]. For classification, we calculated the length of the Rayleigh vector for 5000 shuffled spike trains and compared it to the length of the cell’s Rayleigh vector. Cells were defined as head direction cells when their score were significantly higher than shuffled scores (p<0.05).

### Data analysis of speed cells

To classify speed cells, we calculated the correlation coefficient between the cellular firing rate and the speed of the fish, as is commonly done [45]. Then, 5000 shuffled spike trains were obtained by first calculating the inter-spike intervals, shuffling the inter-spike intervals by random permutation and using the cumulative sum to obtain the shuffled spike train. For each shuffled spike train the correlation coefficient between the firing rate and swimming speed was calculated. Comparing the result of the shuffled spike trains to the cellular result yielded an estimate of the p-value of each cell. The cell was defined as a speed cell if p<0.05.

### Data analysis of velocity cells

To classify velocity cells, we calculated the correlation coefficient between the cellular firing rate and the velocity of the fish. Then, 5000 shuffled spike trains were obtained by first calculating the inter-spike intervals, shuffling the inter-spike intervals by random permutation and using the cumulative sum to obtain a shuffled spike train. For each shuffled spike train, the correlation coefficient between the firing rate and fish’s two-dimensional velocity was calculated. By comparing the result of the shuffled spike trains to the cellular result, we obtained an estimate of the p-value of each cell. A cell was defined as a velocity cell if p<0.05.

## Acknowledgments

We are grateful to Jacob Vecht from Deuteron Technologies Ltd, Nachum Ulanovsky and the members of the Ulanovsky lab for all their help in setting up the recording system for behaving fish. We also thank Nachum Ulanovsky for commenting on the text. In addition, we are grateful to Mario Wullimann for guidance in interpreting the anatomical data and to Tal Novoplansky-Tzur for helpful technical assistance. We gratefully acknowledge financial support from THE ISRAEL SCIENCE FOUNDATION – FIRST Program (grant no. 281/15), the Frankel center at the Computer Science Department, and the Helmsley Charitable Trust through the Agricultural, Biological and Cognitive Robotics Initiative of Ben-Gurion University of the Negev.

## Author Contributions

Conceptualization, E.V., O.B., O.D., R.S., Methodology E.V., L.C., O.B., O.D., R.S., Formal Analysis, E.V., L.C, S.P., O.D., R.S., Investigation E.V., LC, R.S., Writing, E.V., LC, O.B., O.D., R.S.

## Supplementary Material

**Supplementary Figure 1.**
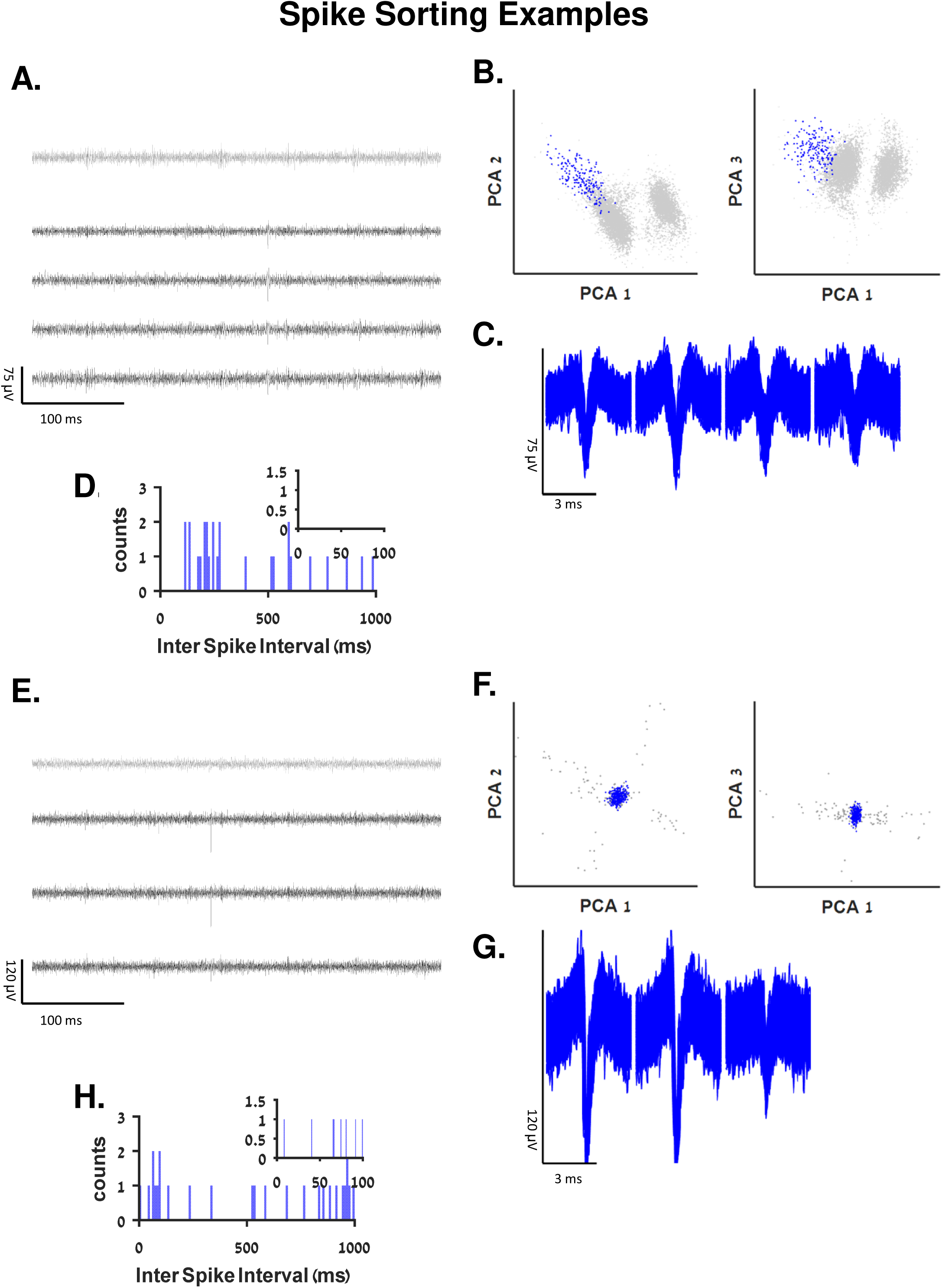
Additional Spike Sorting Examples. **A** and **E.** Examples of raw recordings from a tetrode (black traces) and a reference electrode (gray) in the fish’s lateral pallium. Neural activity can be seen in the tetrode alone. **B** and **F**. Projection on the first three principal components of the data from the tetrode of all candidate spike that crossed the threshold. In panel **F** only one main cluster crossed the detection threshold. **C** and **G.** Waveforms of two neurons after spike sorting. **D** and **H**. Inter spike interval histogram of the detected clusters. Insets show there were no violations of the refractory period in either case. The clusters form a border cell (Figure 2A-D) and a velocity cell (Supplementary Figure 11A-C) respectively.

**Supplementary Figure 2.**
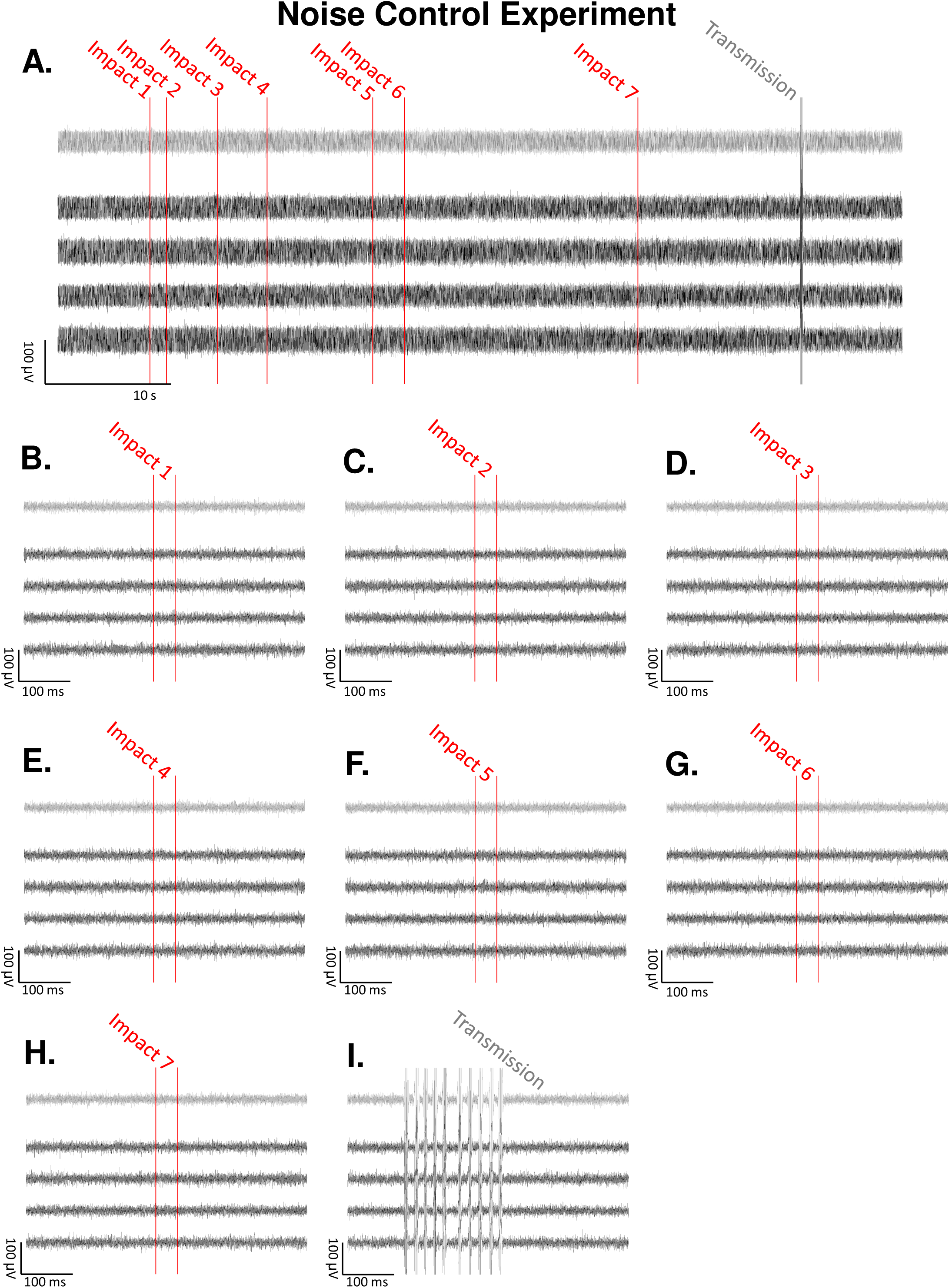
Motion Artifact Control Experiment. To guarantee the absence of spike-like artifacts in the recording system caused by fish motion, for each fish we conducted a control experiment where we generated impacts with the walls while the fish was anesthetized. **A**. Raw recordings from a tetrode (black traces) and a reference electrode (gray) during a control experiment (same recording day and fish as in Figure 1). Impacts with the walls marked by red lines, communication artifact is easily detected. Since the fish was under anesthesia, brain activity was reduced but artifacts could be generated by the impacts. No motion or impact artifacts are present. **B-H.** The start and the end of the frame in which the impact occurred marked in redlines. No motion or impact artifacts are shown. **I.** Communication artifact is easily detected. It affects all of the recording channels and have a unique signature. The logger communicates with the control computer every two minutes in order to synchronize their clocks, the communication occur only every two minutes for about 150 ms.

**Supplementary Figure 3.**
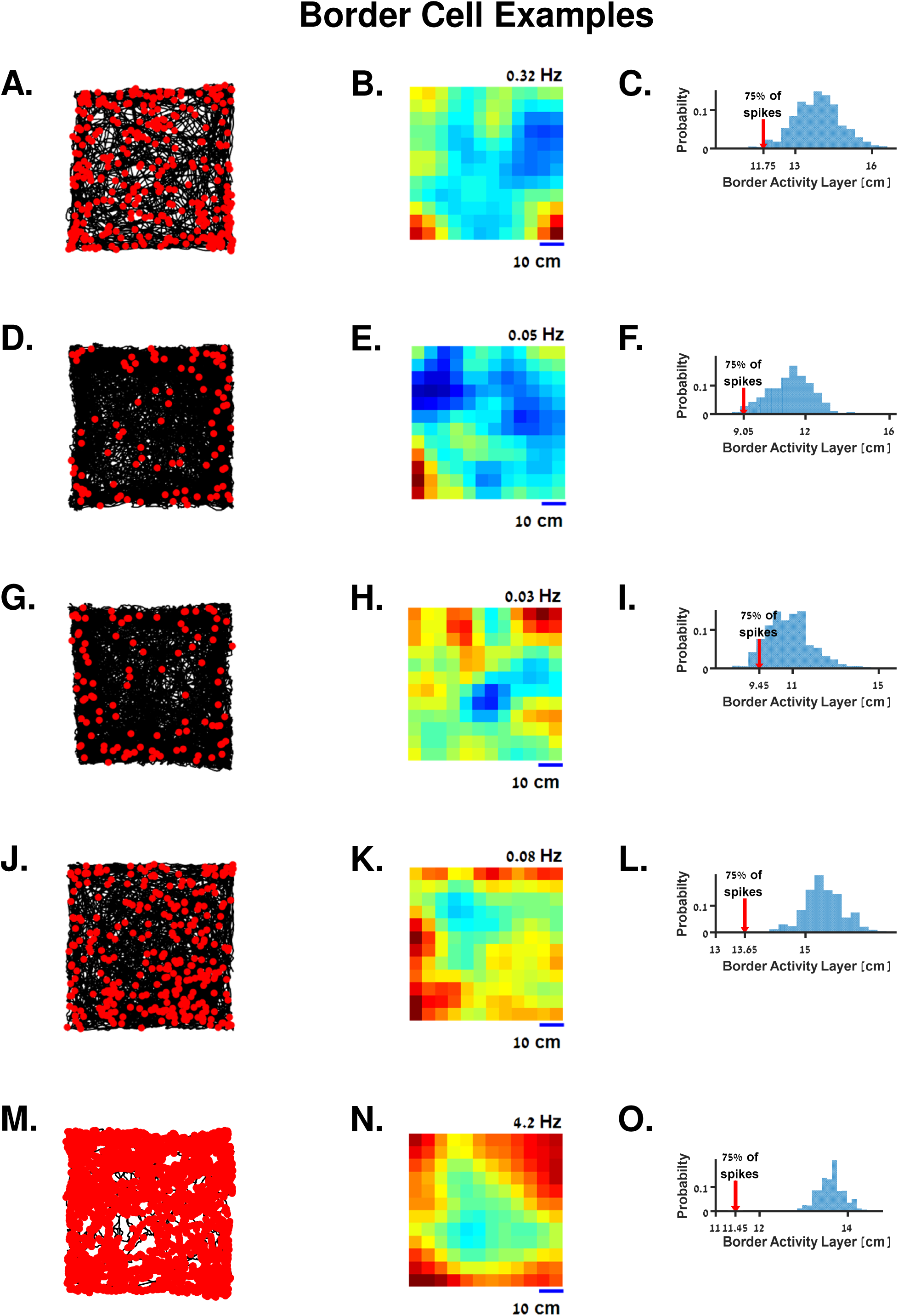
Additional Examples of Border Cells. The cells were recorded in different sessions in different fish. For each cell, the trajectory (black curve, left column) and action potentials (red dots) are presented in addition to a heat map representing the probability of firing in each space bin (middle column) together with a statistical analysis of border activity layer (right column).

**Supplementary Figure 4.**
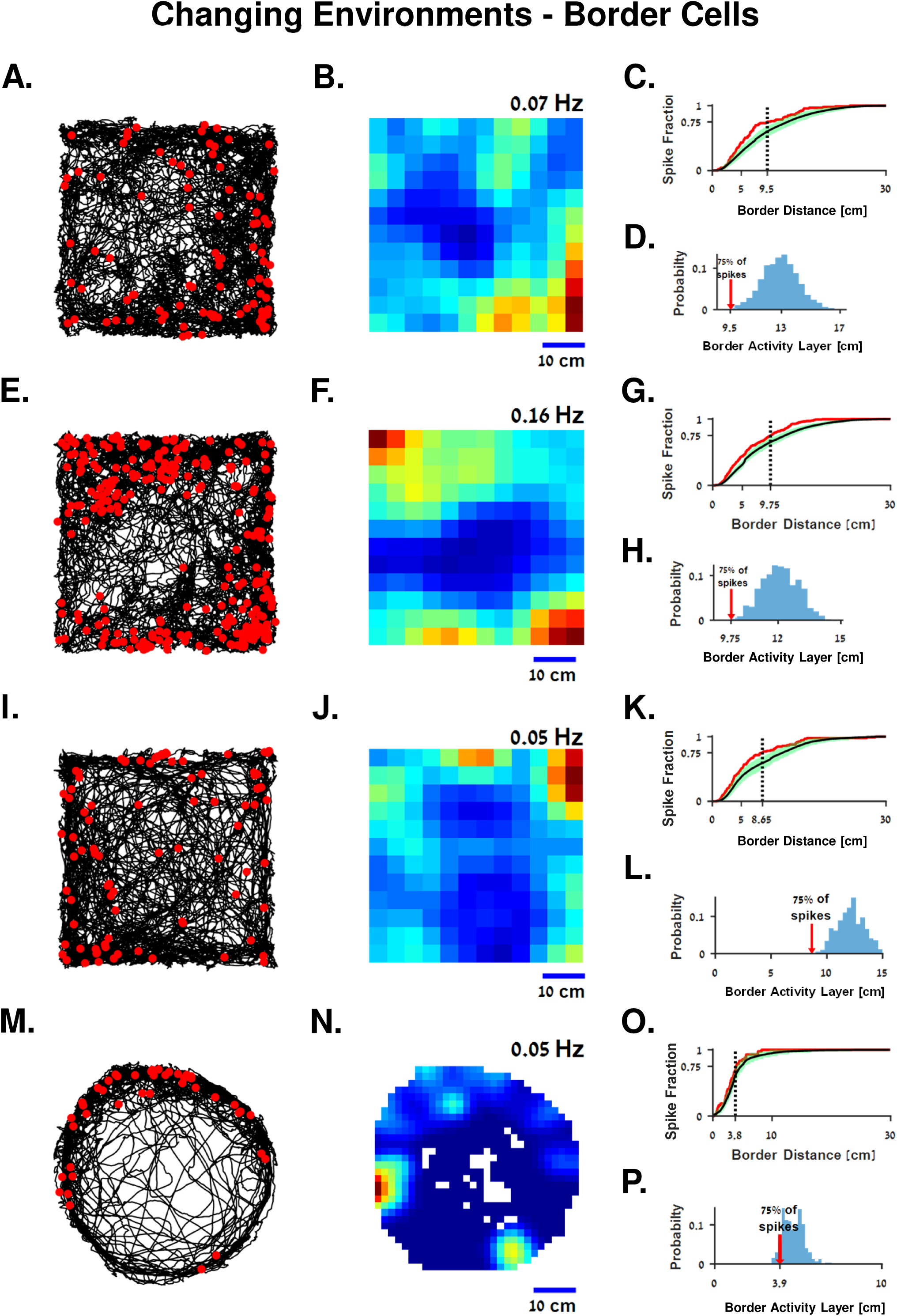
Changing Environments - Border Cells. **A-H** Border cell recorded in two sequential sessions. Before the second session the water tank with visual landmarks marked on it was rotated by 180 degrees. Cell activity shows border proximity characteristics in both cases. A stability test of this cell is shown in Supplementary Figure 4 **I-P** Another border cell recorded in two sequential sessions. Before the second session, the fish was transferred from the square water tank to the circular water tank. Cell activity shows border proximity characteristics in both cases. For each cell, the trajectory (black curve, left column) and action potentials (red dots) are presented in addition to a heat map representing the probability of firing in each space bin (middle column) together with a statistical analysis of border activity layer (right column).

**Supplementary Figure 5.**
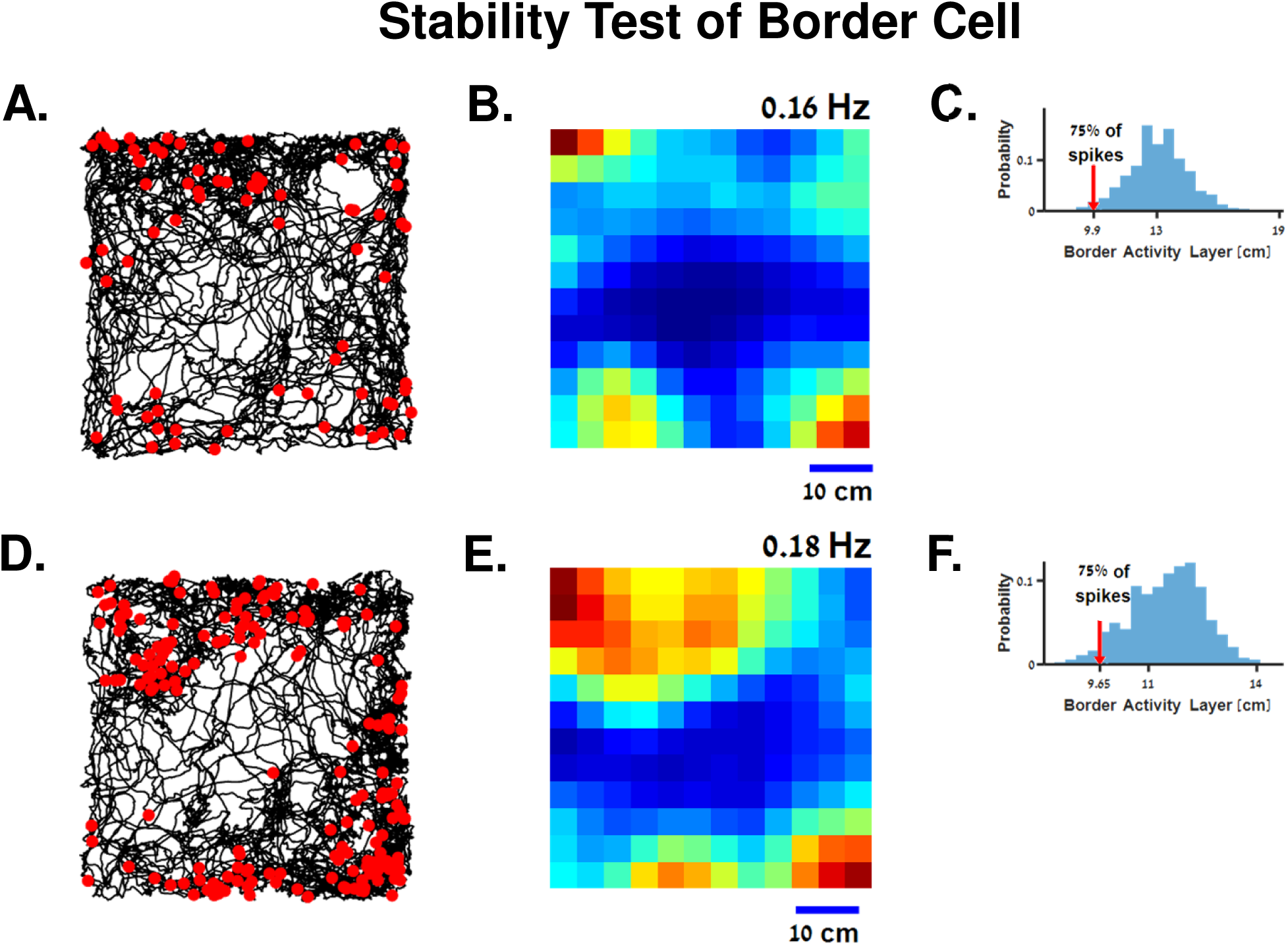
Stability Test of Border Cell. Border cell (same as Supplementary Figure 4 A - H) activity in the first half (A-C) and the second half (D-F) of the same experiment. Cell shows border proximity properties in both parts. The trajectories (black curve, panels A and D) and action potentials (red dots) are presented in addition to a heat map representing the probability of firing in each space bin (panels B and E) together with a statistical analysis of the border activity layer (panels C and F).

**Supplementary Figure 6.**
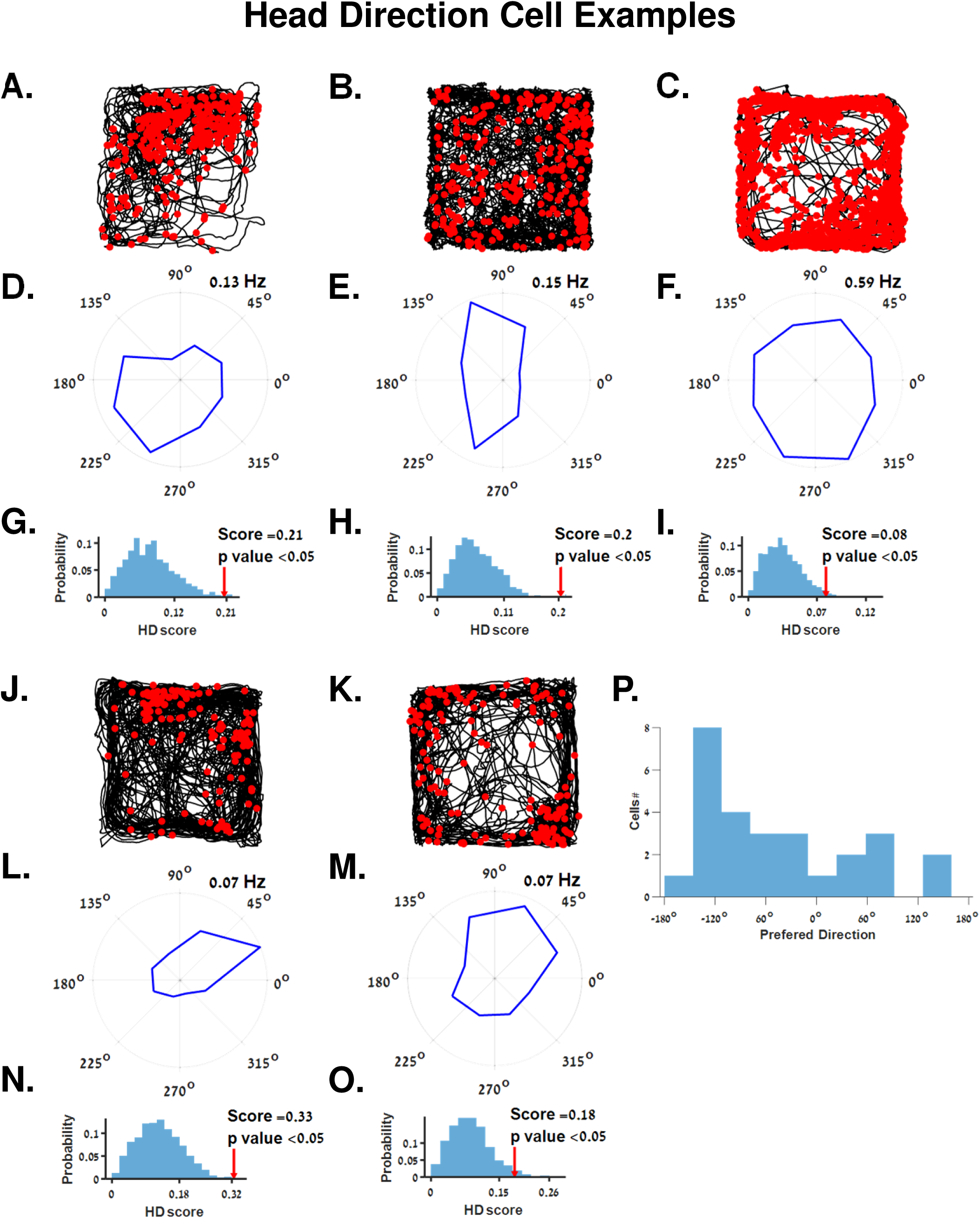
Additional Examples of Head Direction Cells. The cells were recorded in different sessions in different fish. For each cell, the trajectory (black curve panels A, B, C, J and K) and action potentials (red dots) are presented in addition to head direction tuning of the neuron (panels D, E, F, L and M). In addition, the statistical analysis shows that the head direction score of all cells was significant as compared to chance (panels G, H, I, N and O). **P.** Population of preferred direction of all head direction cells.

**Supplementary Figure 7.**
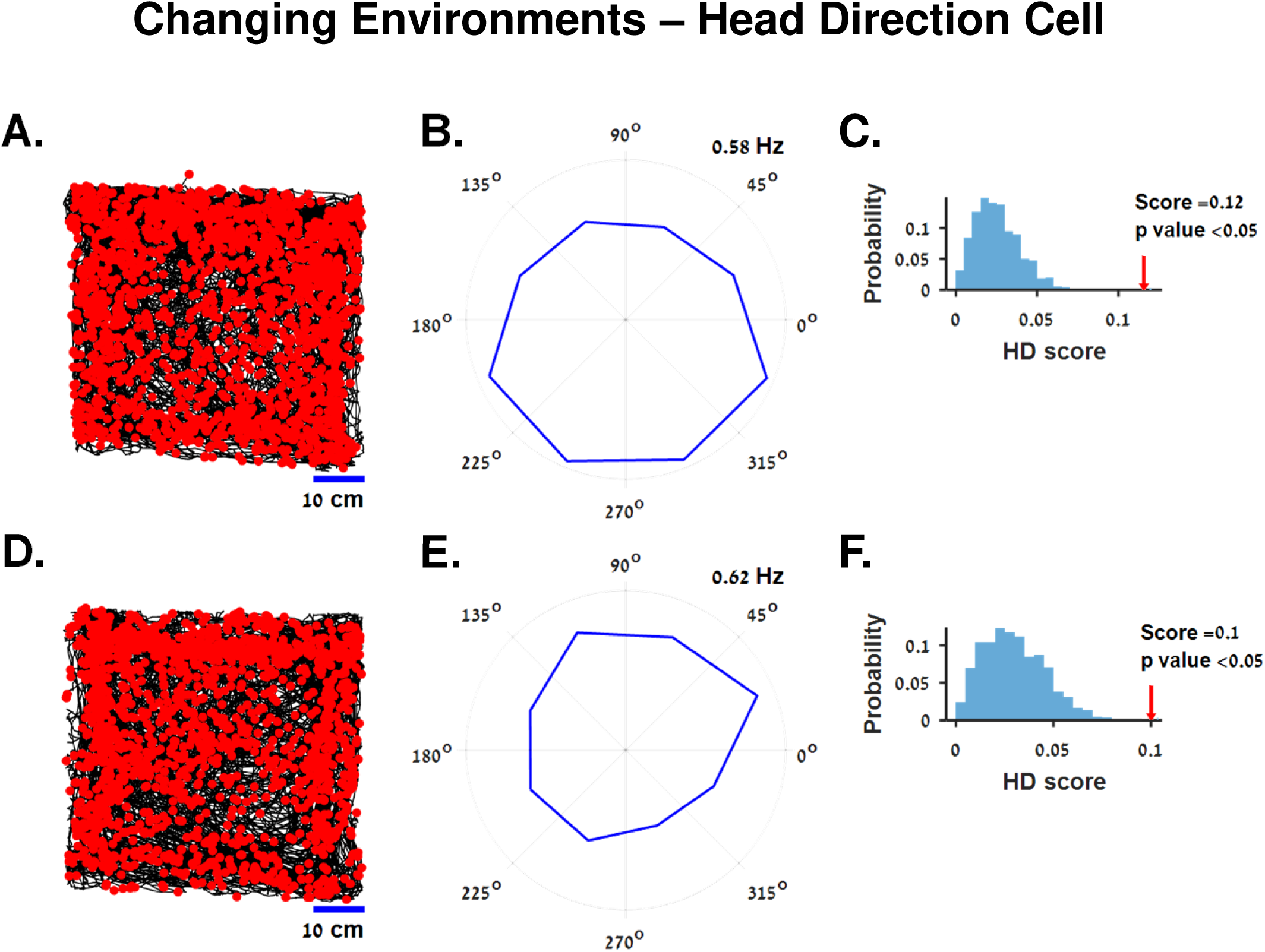
Changing Environments - Head Direction Cell. Head direction cell recorded in two sequential sessions. Before the second session, the water tank with visual landmarks marked on it was rotated by 180 degrees. Cell activity shows a direction preference change corresponding to the rotation.

**Supplementary Figure 8.**
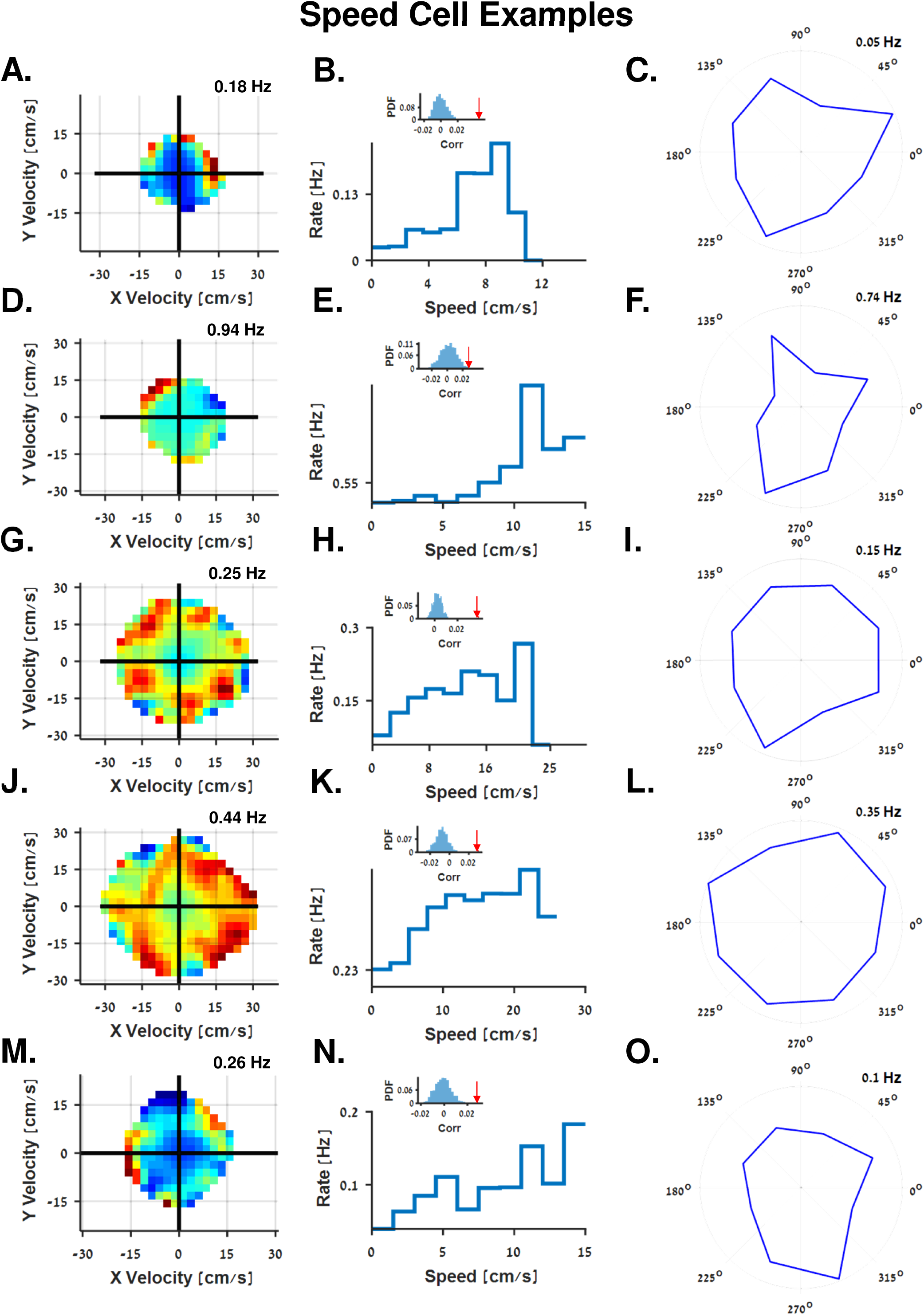
Additional Examples of Speed Cells. The cells were recorded in different sessions in different fish. The dependence of the firing rate on the velocity in two dimensions is presented (left column). Also presented is the rate as a function of speed (middle columns) together with the statistical analysis (middle column, insets), and rate tuning as a function of direction (right column).

**Supplementary Figure 9.**
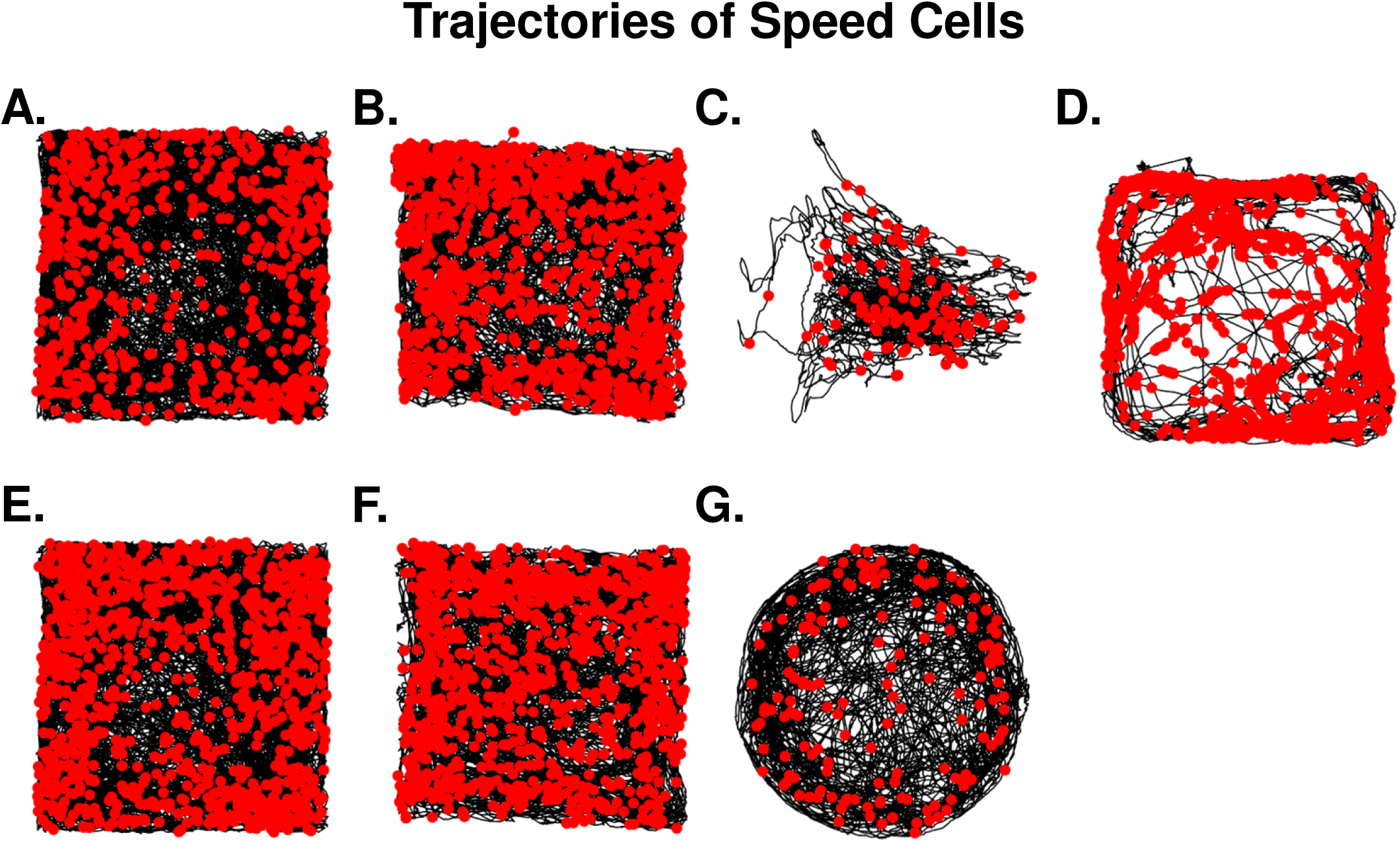
Trajectories of Speed Cells. The fish’s trajectories (black curve) and action potentials (red dots) recorded in all speed cells shown in the previous figures. Panels **A-B** correspond to cells presented in Figure 4 A-F. Panels **C-F** correspond to cells presented in Supplementary Figure 8.

**Supplementary Figure 10.**
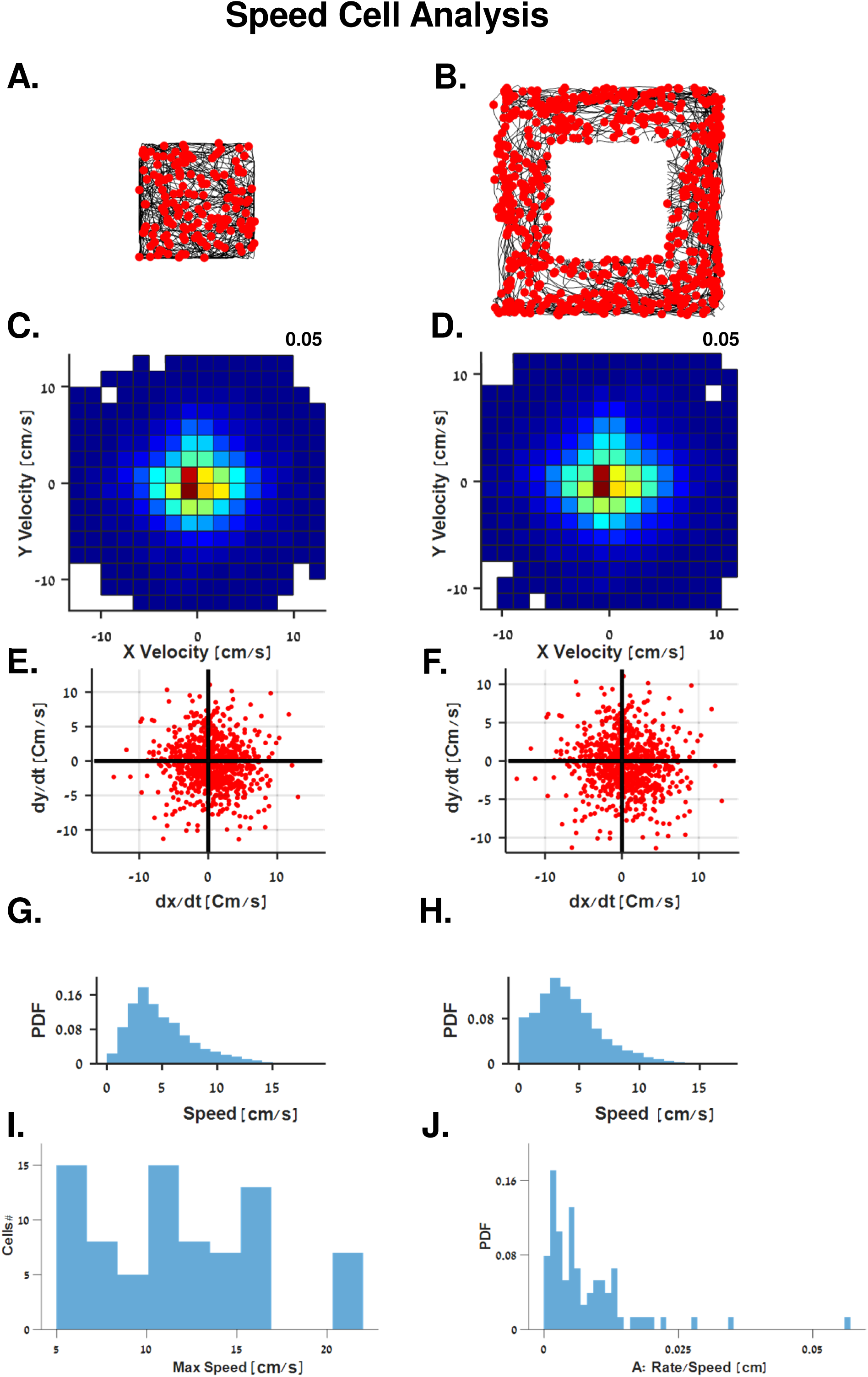
Speed Cell Analysis. **A, C, E and G.** Speed cell activity in the center of the water tank, in comparison to its activity near the aquarium walls (Panels B, D, F and H). The swimming speeds and spiking activity were very similar in both areas of the water tank. **A-B.** Trajectories (black curves) and action potentials (red dots) of the speed cell. **C-D.** Probability density function of swimming velocities, color coded from dark blue (no swimming) to dark red (probability indicated in the upper right side of panels). **E-F.** Fish velocity when action potentials (red dots) occurred. **G-H.** Scalar speed probability density function of the fish. **I.** Population of maximal speeds swam by the fish from all the recorded speed cells. **J**. Probability density function of the speed cells tuning curves’ slopes.

**Supplementary Figure 11.**
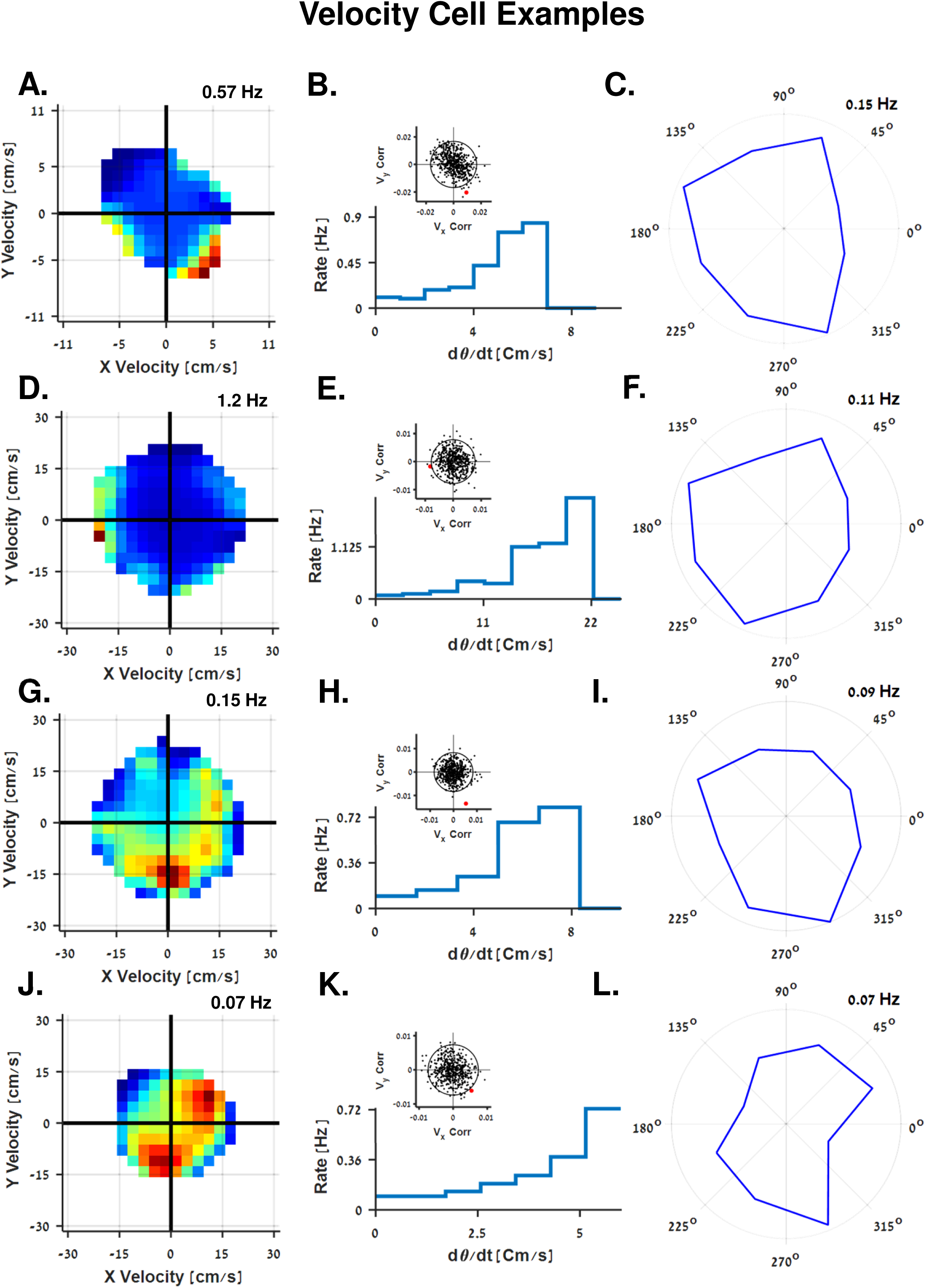
Additional Examples of Conjunction Speed Cells and Head Direction - i.e. velocity cells. Each row of panels corresponds to a single cell. The velocity tuning of the cell can be summarized in the color-coded firing rate map as a function of the two-dimensional velocity (left column). The dependence of the firing rate on the speed along the one-dimensional projections of the preferred direction (middle column) is shown together with a statistical analysis of the velocity tuned cells (middle column, insets): the correlation coefficient between the spike train and the two Cartesian velocity components was calculated for shuffled spike trains (black dots) and the data (red dots). The 95% confidence circle is depicted. Right column: directional tuning of each cell. **A - C**. Waveform of the cell spiking activity is presented in Supplementary Figure 1G

**Supplementary Figure 12.**
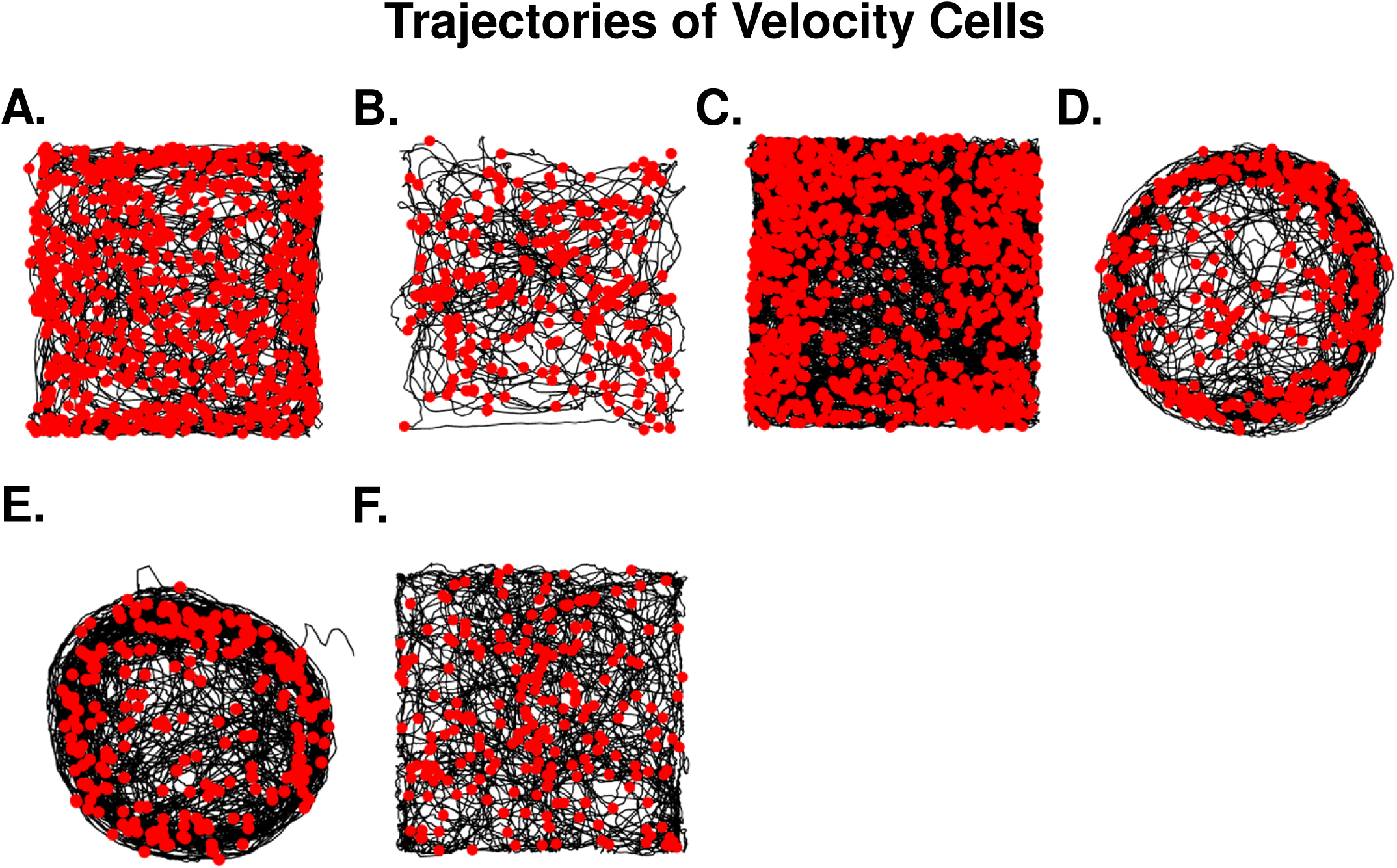
Trajectories of Velocity Cells. The fish’s trajectories (black curve) and action potentials (red dots) recorded in all velocity cells shown in previous figures. Panels **A-B** correspond to cells presented in Figure 4 G-L. Panels **C-F** correspond to cells presented in Supplementary Figure 11.

**Supplementary Figure 13.**
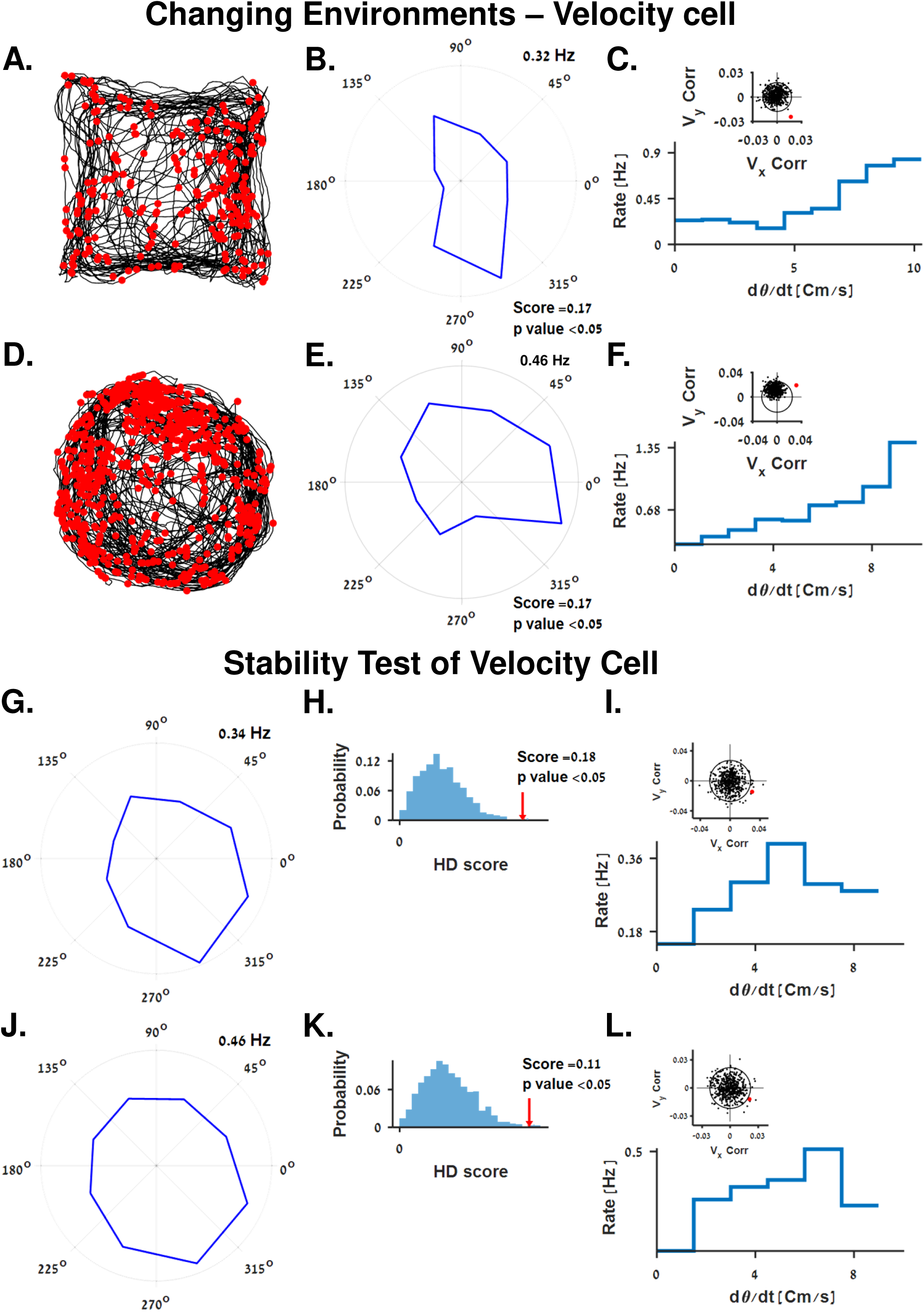
Changing Environments - Velocity Cell, Stability Test of Velocity Cell. **A-F.** Velocity cell recorded in two sequential sessions. Before the second session the fish was transferred from the square water tank to the circular water tank. Cell activity shows both head direction preference and a correlation to velocity in both cases. Waveform of the cell spiking activity is presented in Figure 1D (blue cluster) **A** and **D**. Trajectories (black curves) and action potentials (red dots) of the presented velocity cell. **B** and **E**. Head direction tuning of the neuron. Head direction score and p value compared to chance are indicated on the bottom right side of panels. **C** and **F.** Firing rate tuning curve to velocities of velocity cells. Insets show the statistical analysis of the velocity tuned cells. **G-L** velocity cell’s activity in the first half (G-I) and the second half (J-L) of the same experiment. Cell shows both head direction preference and a correlation to velocity in both parts. The head direction tuning of the neurons is presented (panels G and J). Statistical analysis of head direction scores (red arrows) shows significantly higher values than chance (panels H and K). **I and L.** Firing rate tuning curve to velocity in each half of the experiment. Insets show statistical analysis of the velocity tuned cell.

**Supplementary Figure 14.**
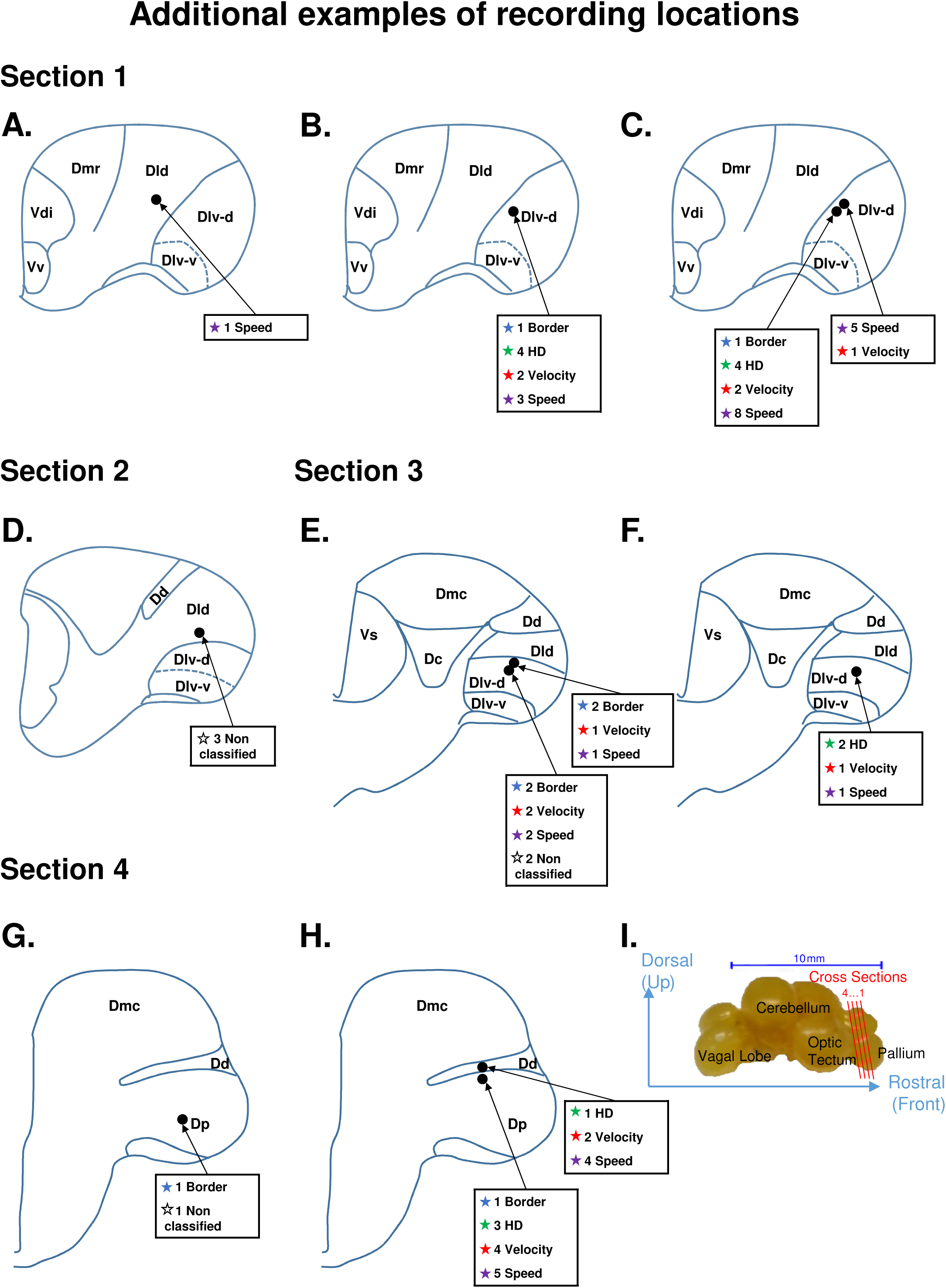
Additional examples of recording locations. **A-H.** Recording locations together with the classification of cells found within this location from 8 different fish. **I.** Goldfish brain structure. Cross sections 1-4 correspond to the section numbers indicated above panels A, D, E and G.

**Supplementary Figure 15.**
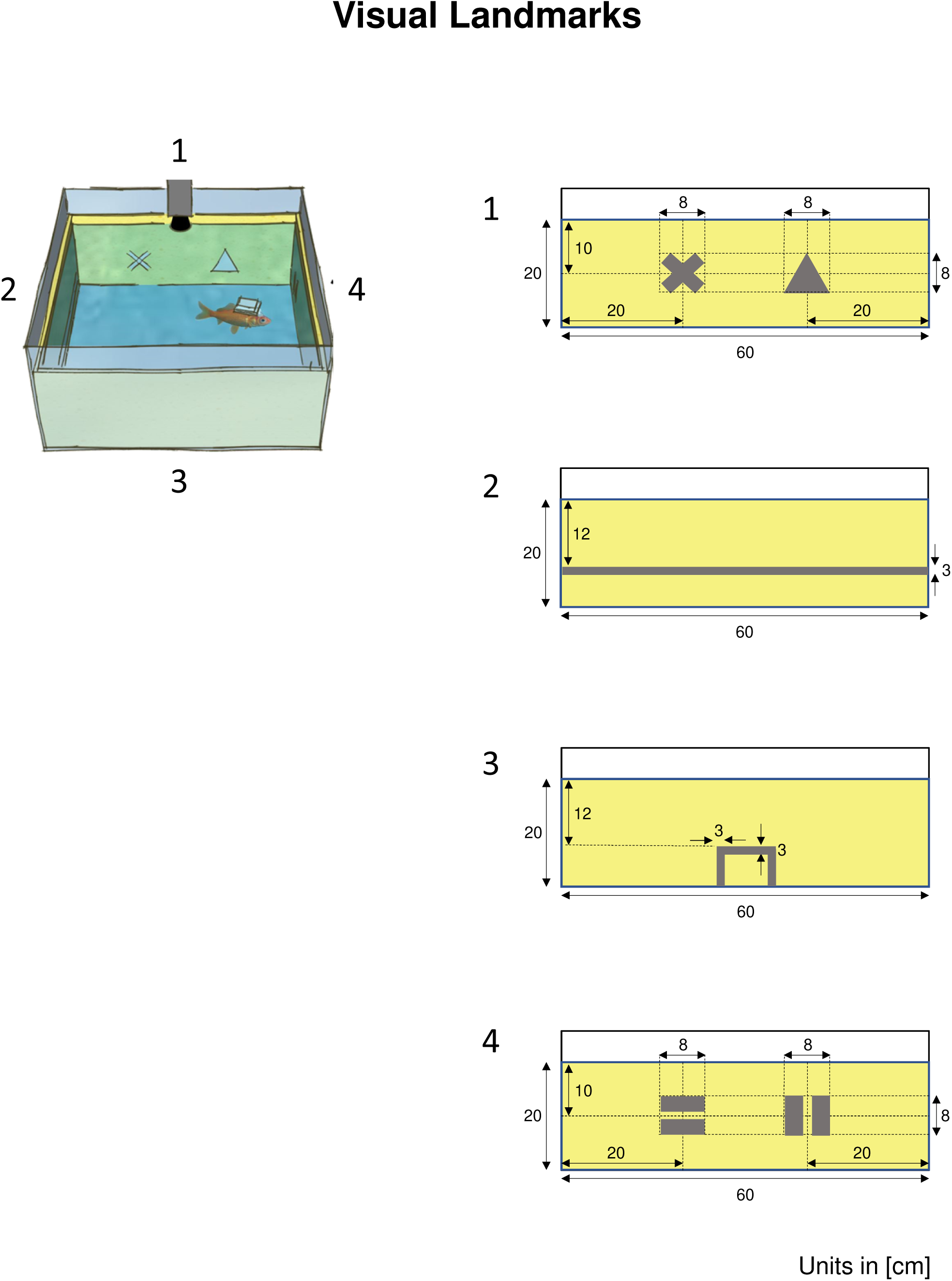
Visual Landmarks. Schematic overview of the experimental water tank and visual cues painted on its walls.

